# Vascular niches are the primary hotspots for aging within the multicellular architecture of cardiac tissue

**DOI:** 10.1101/2025.03.30.646237

**Authors:** David Rodriguez Morales, Veronica Larcher, Mariano Ruz Jurado, Lukas Tombor, Lukas Zanders, Julian U. G. Wagner, Andreas M. Zeiher, Christoph Kuppe, David John, Marcel Schulz, Stefanie Dimmeler

**Affiliations:** Goethe University Frankfurt, Institute for Cardiovascular Regeneration, Theodor-Stern-Kai 7, 60590 Frankfurt am Main, Germany; German Centre for Cardiovascular Research (DZHK), Frankfurt am Main, Germany; Cardiopulmonary Institute, Goethe University Frankfurt am Main, Germany; Division of Nephology and Clinical Immunology, RWTH Aachen University, Medical Faculty, Aachen, Germany; Goethe University Frankfurt, Institute for Computational Genomic Medicine, Theodor-Stern-Kai 7, 60590 Frankfurt am Main, Germany

**Keywords:** Aging, Senescence, Spatial-Transcriptomics

## Abstract

**Background:** Aging is a major, yet unmodifiable risk factor for cardiovascular diseases, leading to vascular alterations, increased cardiac fibrosis, and inflammation, all of which contribute to impaired cardiac function. However, the microenvironment inciting age-related alterations withing the multicellular architecture of the cardiac tissue is unknown.

**Methods:** We investigated local microenvironments in aged mice hearts applying an integrative approach combining single-nucleus RNA sequencing and spatial transcriptomics in 12-week-old and 18-month-old mice. We defined distinct cardiac niches and studied changes in their cellular composition and functional characteristics.

**Results:** Integration of spatial transcriptomics data across young and aged hearts allowed us to identify 11 cardiac niches, which were characterized by distinct cellular composition and functional signatures. Aging did not alter the overall proportions of cardiac niches but leads to distinct regional changes, particularly in the left ventricle. Whereas cardiomyocyte-enriched niches show disrupted circadian clock gene expression, vascular niches showed major changes in pro-inflammatory and pro-fibrotic signatures and altered cellular composition. We particularly identified larger vessel-associated cellular niches as key hotspots for activated fibroblasts and macrophages in aged hearts, with interactions of both cell types through the C3:C3ar1 axis. These niches were also enriched in senescence cells exhibiting high expression of immune evasion mechanisms that may impair senescent cell clearance.

**Conclusion:** Our findings indicate that the microenvironment around the vasculature is particularly susceptible to age-related changes and serves as a primary site for inflammation-driven aging, so called “inflammaging”. This study provides new insights into how aging reshapes cardiac cellular architecture, highlighting vessel-associated niches as potential therapeutic targets for age-related cardiac dysfunction.

## Introduction

Aging is a natural physiological process associated with structural and functional changes and cellular senescence that can impair cardiovascular health. These alterations significantly increase the risk of developing cardiovascular diseases, the leading cause of death worldwide^1^. The aging heart is characterized by increased fibrosis, subsequent myocardial stiffening and reduced cardiac compliance^2^. Diastolic dysfunction becomes more pronounced, resulting in elevated filling pressures. Additionally, both microvascular and macrovascular function deteriorate with age, contributing to endothelial dysfunction, reduced perfusion, and increased vascular stiffness. Aging also affects cardiac immune cells, particularly inflammatory macrophages (MP), shifting them towards a more pro-inflammatory state^2^. Particularly, Ccr2^+^ bone marrow-derived MP increase, driving chronic low-grade inflammation. This pro-inflammatory environment, known as “inflammaging”, plays a key role in cardiac senescence by promoting oxidative stress, DNA damage, and the activation of senescence-related pathways. These immune and structural changes accelerate cardiovascular disease progression and increase the risk of heart failure with preserved ejection fraction in older individuals.

In recent years, advancements in omics technologies, particularly single-cell RNA-sequencing (scRNA-seq), have revolutionized the study of cardiac biology. These techniques provide an unbiased framework for examining cellular heterogeneity in the heart^3,4^; enabling the precise characterization of gene expression profiles at the single-cell resolution. However, the information generated lacks the spatial context. Several platforms have been developed to perform spatial transcriptomics^5^, allowing to study changes in the spatial dynamics of the heart^6,7^.

To explore the spatial organization and molecular changes in the aging heart on a whole-transcriptome level, we generated spatial transcriptomics data obtained through the 10X Genomics Visium Platform from 12-week and 18-month-old murine hearts. Integration of this data with single-nucleus RNA sequencing (snRNA-seq) enabled the characterization of cardiac niches and the assessment of changes in cell composition and interactions. Notably, vascular niches emerged as key regions of aging-associated alterations, showing increased expression of fibrosis-related genes and a shift in macrophage populations, fostering a pro-fibrotic and pro-inflammatory environment. These niches also served as primary hotspots for cellular senescence in the aging heart.

## Methods

### Animals

3- and 18/20-month-old female and male C57Bl/6J wildtype mice purchased from Janvier (Le Genest Sain-Isle, France) were used in this study.

### Mouse tissue processing

Heart tissues for the spatial transcriptomics assay were immediately frozen in liquid nitrogen after OCT compound (Tissue-Tek) embedding. Ten-micrometer tissue cryosections were stained with hematoxylin and eosin (H&E) and the appropriate tissue regions were selected for further processing.

### Spatial transcriptomic gene expression assay

For the single spatial transcriptomics assay, frozen heart tissue from 3-month (n=5) and 18-month-old (n=5) female mice were embedded in OCT compound (Tissue-Tek) and subsequently cryosectioned at 10 µm thickness using a Cryostar NX70 cryostat (Thermo Fisher Scientific). Tissue sections were immediately transferred onto pre-chilled Visium Tissue Optimization Slides (10X Genomics, PN-1000193) to determine the optimal permeabilization time. Following manufacturer recommendations (10X Genomics), optimization experiments established an ideal permeabilization window at either 12 or 18 minutes for efficient tissue digestion and RNA release.

For spatial transcriptomics, tissue sections were placed onto Visium Spatial Gene Expression Slides (10X Genomics, PN-1000187) and processed according to the Visium Spatial Gene Expression User Guide. H&E staining was performed as suggested by 10X Genomics. Brightfield imaging of tissue sections was performed using a Fritz brightfield digital microscope (Precipoint). Image stitching and processing were conducted using NIS-Elements software (Nikon).

Next-generation sequencing libraries were prepared following standard protocols provided in the Visium User Guide. Libraries were loaded at a concentration of 300 pM and sequenced on an Illumina NovaSeq 6000 System using parameters recommended by 10X Genomics.

### Immunohistochemistry

Hearts from 3-month (n=6), 18 to 20-month (n=5) female and male mice were perfused and fixed using 4% PFA (28908, Thermo Fisher Scientific) overnight, followed by rinsing three times with PBS. Tissues were dehydrated in a sucrose gradient (5% for 2 hours, 12% for 2 hours, 20% overnight) in a 15% sucrose, 8% gelatine (G1890, Sigma-Aldrich) and 1% polyvinylpyrrolidone (PVP, P5288, Sigma-Aldrich) solution. 50 μm sections were cut using a Leica cryostat (CM3050 S), mounted on Superfrost Plus microscope slides (10149870, Fisher Scientific), and stored at −20°C (short-term) or −80 °C (long-term) until being processed for immunofluorescence. For staining, slices were thawed and rehydrated by washing with PBS for 5 min. Slices were permeabilized for 30 minutes in PBS-0.5% Triton X-100 and blocked with 10% normal donkey serum (AB7475, Abcam), 3% bovine serum albumin (BSA, A7030, Sigma-Aldrich), 0.1% Triton X-100 in PBS for 1 hour at room temperature. Primary antibodies were incubated at 4 °C overnight. The following primary antibodies were used: anti-Pdgfra (AF1062, R&D Systems), anti-Smooth Muscle Actin (5694, Abcam), anti-NG2 (AB5320, Millipore), anti-Cd68 (97778S, CellSignalling), anti-Lyve1 (AF2125, R&D Systems), Isolectin B4 biotinylated (B-1205, Vector). Sections were washed three times with PBS-0.1% Triton X-100 and incubated with secondary antibodies for 1 hour at room temperature. The following secondary antibodies were used: donkey anti-rabbit Alexa Fluor 647 (A31573, Thermo Fisher Scientific), donkey anti-rabbit Alexa Fluor 555 (A31572, Thermo Fisher Scientific), donkey anti-goat Alexa Fluor 488 (A11055, Thermo Fisher Scientific), donkey anti-goat Alexa Fluor 647 (A21447, Thermo Fisher Scientific), Streptavidin Alexa Fluor 555 conjugate (S32355, Invitrogen). Sections were washed three times with PBS-0.1% Triton X-100, counter-stained with DAPI and mounted with Dako Fluorescence Mounting Medium (S3023, Agilent Dako). Immunofluorescence images were acquired using a Leica Stellaris confocal microscope.

#### Beta galactosidase staining

Senescence-associated β-galactosidase was performed on cryopreserved sections using the CellEvent Senescence Green Detection Kit (C10850, ThermoFisher) according to manufacturer’s instruction. Briefly, slices were thawed and rehydrated by washing with PBS for 5 min. CellEvent Senescence green probe were diluted into pre-warmed CellEvent Senescence Buffer and incubated at 37 °C for 4 hours.

### Quality control and processing of the transcriptomics data

Data processing of all the transcriptomics data was performed using Scanpy (v1.10.1)^8^, operated under Python v3.11.8, and Seurat (v5.0.1)^9^, operated under R v4.4.1.

### Single-nuclei RNA sequencing processing

The snRNA-seq FASTQ files were mapped to the mouse reference genome (GRCm38) as previously described^10,11^. The count matrix generated by the alignment tool of the individual samples was combined into one AnnData object and subjected to a filtering process. Nuclei with a low number of genes (< 250), and an excess of mitochondrial RNA (> 5%), were excluded. Doublets were identified and removed using Scanpy’s wrapper of scrublet^12^ with default parameters. Genes expressed in less than 3 nuclei were not considered for further analysis. Gene expression values were normalized and logarithmize using a scaling factor of 10,000 using the sc.pp.normalize_total and sc.pp.log1p functions.

Highly variable genes (HVGs) were identified using the *sc.pp.highly_variable_genes* function with the default arguments and setting the *batch_key* argument to identify HVGs shared across samples. The HVGs were used for the integration using single-cell Variational Inference (scVI)^13^, a method based on a conditional variational autoencoder available in the scvi-tools package (v1.0.4, *n_hidden=128*, *n_latent=50*, *n_layers=3*, *dispersion=’gene-batch’*), which was configurated to account for batch effects, including 10X Genomics library generation kit differences. We accounted for unwanted sources of variation (total counts, and percentage of mitochondrial, and ribosomal genes). The latent representation of the autoencoder was used to estimate the nearest neighbors’ distance matrix using the BBKNN package (v1.6.0, *neighbors_within_batch=8*)^14^. Scanpy functions were then used to estimate the uniform manifold approximation and projection (UMAP) embeddings (*sc.tl.umap*, *spread=1.2*, *min_dist=.3*) and cluster the data using the Leiden algorithm (*sc.tl.leiden*, *resolution=5 and resolution=1.5*). Clusters were annotated automatically using celltypist (v1.6.3, Human Heart model v1.0)^15^ and manually cross-checked with gene markers^16,17^. A cluster was annotated as neural cells; however, marker expression did not support the annotation and therefore was excluded from further analysis. Re-clustering of endothelial cells and myeloid cells was performed by applying BBKNN (*neighbors_within_batch=3*) and the Leiden algorithm (*resolution=1.5*). Differential gene expression analysis was performed using the *sc.tl.rank_genes_groups* (*method=’wilcoxon’, tie_correct=True*) function from Scanpy. Differential gene expression (DGE) analysis between young and old condition was performed considering only the cells derived from Vidal et al^10^. The cells belonging to the Wagner et al^11^ dataset was excluded from the DGE analysis due to the usage of a different 10X library kit and the inclusion of treatment conditions.

### Spatial transcriptomics processing

The spatial transcriptomics FASTQ files and histology images were processed and mapped to the mouse reference genome (GRCm38, “refdata-gex-mm10-2020-A”) using the SpaceRanger software (v2.0.1) using the default parameters. A permissive filtering was applied to the count matrices. We remove spots with a low number of UMI counts (< 500) and genes (< 300) as well as genes lowly expressed (< 3 spots). Mitochondrial and hemoglobin genes were excluded from the analysis, as they do not provide relevant spatial information. Normalization was performed using the *SCTransform* function from Seurat (*vst.flavor=“v2”* and *min_cells=3*). The raw counts were also normalized and logarithmize using a scaling factor of 10,000 using the sc.pp.normalize_total and sc.pp.log1p functions from Scanpy. Samples were integrated to identify shared spatial domains using STAligner^18^. First, a spatial neighbor network was constructed for each section using *Cal_Spatial_Net* (*radius=125*), and then the HVGs shared across the different sections were selected for training a graph attention auto-encoder to generate a spatially aware embedding using *train_STAligner* (*hidden_dims = [450, 30]*, *knn_neigh=100*, *margin=2.5*). The latent representation of the autoencoder was used to estimate the nearest neighbors distance matrix using BBKNN (*neigbors_within_batch=5*), and Scanpy functions were used to estimate the UMAP embeddings (*sc.tl.umap*, *spread=1*, *min_dist=0.2*), and cluster the data using the Leiden algorithm (*sc.tl.leiden*, *resolution=0.8)*.

The pseudo-bulk expression matrix was obtained using the Python implementation from decoupleR^19^ (v1.8.0), a package containing different statistical methods to extract biological activities from omics data. Raw UMI counts were summed using the *get_pseudobulk* (*min_cells=10*, *min_counts=1000*). Subsequently, the matrix was log-normalized using a scaling factor of 10,000 as previously described, and principal component analysis was performed. Associations between the metadata and the principal components were estimated using ANOVA, implemented in the *get_metadata_associations* function. The default arguments were used for the estimation, including a significance threshold of 0.05 and the usage of the Benjamini-Hochberg method for multiple testing correction.

### Deconvolution of spots

Spatial transcriptomics sections were processed to estimate the cell-type composition for each spot using cell2location (v0.1.3)^20^ under Python v3.9.18. The integrated and annotated snRNA-seq gene expression values were pre-filtered using *filter_genes* (*cell_count_cutoff=3*, *cell_percentage_cutoff2=0.03*, *nonz_mean_cutoff=1.12*). Estimation of the reference cell type signatures was performed using regularized negative binomial regressions accounting for technical effects (10X Genomics library generation kit differences) and batch effects. Afterwards, the sections were divided into two groups (young and old) and were deconvoluted simultaneously using a hierarchical Bayesian model as implemented in *cell2location.models.Cell2location* accounting for batch effects. Two hyperparameters need to be provided to the pipeline, the detection alpha, which affects the regularization used for variations in the detection of RNA within experiments, and the expected average number of nuclei per spot. For the young sample, a detection alpha of 200 was used, while 20 was used for the old samples. In both groups, the hematoxylin and eosin staining were used to determine the average number of nuclei per spot (n = 8) and was used as the hyperparameter. After training, the predicted cell type abundances were used to estimate the cell type proportions per spot.

### Annotation of anatomic regions in the heart

Cloupe files generated by SpaceRanger were used to annotate the anatomic regions using the Loupe Browser (v8.0.0) software from 10x Genomics. Three major regions were annotated: the right ventricle, the left ventricle, and the septum. The left ventricle was further divided into 3 sections with similar size (inner, middle, and outer regions). Regions that could not be distinguished correctly were annotated as background and excluded from the analysis.

### Inference of pathway activities and gene set overrepresentation analysis

Inference of pathway activities was performed with the Python implementation of decoupleR^19^, and the PROGENy model^21^, a resource containing a curated collection of pathways, their target genes, and weights for each interaction. The inference was performed using the multivariate linear model implemented in the *run_mlm* function. The median estimated enrichment scores across all samples were used to classify the pathways as activated or inactivated. Pathways with a positive median score were considered to be activated and were further explored by performing differential analysis using the *sc.tl.rank_genes_groups* (*method=’wilcoxon’, tie_correct=True*) function from Scanpy.

In addition to the analysis of pathway activities, we performed an overrepresentation analysis of biological processes from the MSigDB^22,23^ using the *run_ora* function. The inferred enrichment scores were used to perform differential analysis between each cardiac niche and the rest of the niches. Significantly regulated pathways (adjusted p-value < 0.01) were sorted based on the scores obtained from the *sc.tl.rank_genes_groups* function and the top 20 most important processes were visualized with Cytoscape (v3.10.3). Uninformative pathways were excluded from the visualization and similar pathways were grouped. Functional enrichment analysis for each niche across conditions was also performed employing a pseudo-bulk approach. In short, a pseudo-bulk count matrix was generated as discussed in the spatial transcriptomics processing section, and the DESeq2 python wrapper (v0.4.11)^24^ was used to perform differential expression analysis. As suggested by the authors, the t-values, which account for both the direction and significance were used to perform overrepresentation analysis using the *get_ora_df* function.

### Analysis of spatial cell dependencies

We used Misty’^25^ implementation in mistyR (v1.12.0) to estimate the importance of the distribution of one cell type to explain the abundance of the rest of the cell types. It is important to note, that the importance can represent both co-localization and co-exclusion of two pairs of cell types. We analyzed the relationship across cell types defining three different spatial contexts. First, we defined an intra-view or same-spot interaction that explains the relationships within spots, second a juxta-view or hexamer space interaction that captures the abundances of the immediate neighbors (radius of ∼100 µm), and lastly, a para-view or extended tissue interaction which captures the abundance in a larger context (radius of ∼200 µm). To avoid ambiguity, the cell abundance in the juxta-view was not considered in the para-view. After the estimation of the aggregated importance for each cell type pair in each sample, we excluded pairs with negative importance and pairs with a spatial variance signature (R^2^) value below 5. Then, the median importance for each cell type pair was estimated across all samples.

### Inference of cell communication events

Inference of cell communication events (CE) was performed with Holonet^26^ (v0.1.0), a method that models CE in spatial transcriptomics data as a multi-view network based on the expression of the ligand and receptor pairs. In short, for each ligand-receptor pair a graph is generated, where each node represents a single spot, and the edges connect sender and receiver spots. Each edge is weighted based on the expression of the ligand, the receptor, and the distance between nodes, and afterward, edges with low confidence are removed. Spatial transcriptomics sections were processed separately, and the communication strength was estimated considering a short diffusion distance (maximum 100 µm). Only ligands and receptors expressed in at least 5 % of the spots were included in the analysis. To visualize the CE network, this method defines CE hotspots based on eigenvector centralities that allow the visualization of regions with activated communication.

### Identification and definition of hotspots of senescence

We used *sc.tl.score_genes* function from Scanpy to score each spot in the spatial transcriptomics for the expression of genes in a gene set. This function reproduces the approach in Seurat^27,28^ by computing a score for each spot that is equal to the mean expression of genes in each spot minus the average expression of genes in a background gene set. The background gene set is composed by randomly selecting 50 genes with similar average expression (computed over all spots). We define a senescence signature gene set by combining five previously published gene sets: 1. CellAge^29^ (370 genes), 2. Aging Atlas^30^ (391 genes), 3. SenMayo^31^ (118 genes), 4. Cellular Senescence from MSigDB^23^ (141 genes), and 5. Fridman and Tainsky^32^ (101 genes)). In addition, only differentially expressed genes (adjusted p-value < 0.05) obtained from comparing young and old sections were used, generating a gene set comprising 593 genes. After scoring all spots based on our gene set, we ranked the spots based on their senescence score over all the samples to avoid biases due to a differential number of spots between samples. The top 5 % of spots with the highest senescence score were annotated as hotspots of senescence (Hspot). Next, we annotated the surrounding spots around the Hspots based on their distance (100, 200, 300, 400, 500 and >500 µm).

### Data availability

The raw data and the AnnData object of the spatial transcriptomics data is available at Biostudies under the accession number E-MTAB14947. The AnnData object with the communication event analysis of the spatial transcriptomics is available at Figshare (https://figshare.com/s/4cbb58c8d22c468d7488). The raw data of the previously published snRNA-seq is available in ArrayExpress under the accession number E-MATB13093 and E-MATB7869. The processed AnnData object generated in this paper is available at Figshare (https://figshare.com/s/c7495368602390abc69b).

### Code availability

All code used for the analysis is available at https://github.com/davidrm-bio/RodriguezMorales_et_al_2025.

## Results

### Multi-omics integrative analysis for the characterization of the heart

We applied an integrative approach to explore and characterize the cardiac changes upon aging using previously published murine snRNA-seq^10,11^ and newly generated spatial transcriptomics data obtained through the 10X Genomics Visium Platform (**Figure 1a, b**).

**Figure 1.**
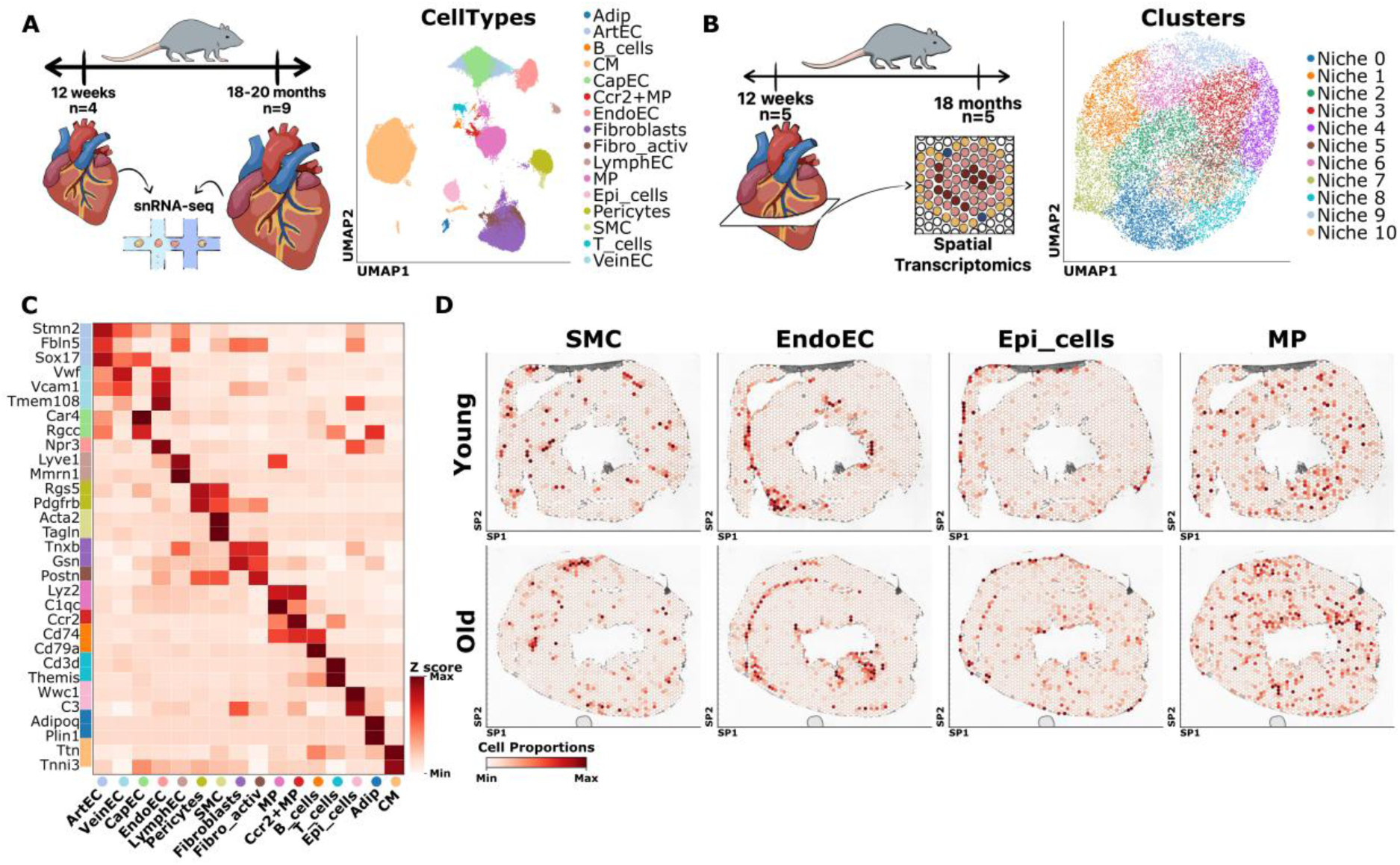
Multi-omics characterization of the aging heart in mice. A) and B) Study overview and UMAPs of all the samples of the cell type annotation in the snRNA-seq (n=64,845) and the niche annotation in the spatial transcriptomics data (n=15,838). Arterial endothelial cells (ArtEC); venous endothelial cells (VeinEC); capillary endothelial cells (CapEC); endocardial endothelial cells (EndoEC); lymphatic endothelial cells (LymphEC); Smooth muscle cells (SMC); Macrophages (MP); Epicardial cells (Epi_cells); and Cardiomyocytes (CM). C) Average expression of marker genes in the snRNA-seq after Z-score transformation. The colors assigned to each gene indicate the cell type it identifies and matches the colors in A). D) Examples of the distribution of cell type proportions estimated in the spatial transcriptomics data. The minimum and maximum values were estimated per panel. The maximum color was set to the percentile 99.2 of the cell proportion per panel.

The snRNA-seq consisted of 13 samples from the hearts of 12-week (n=4) and 18 to 20-month-old mice (n=9). After quality control, 64,845 nuclei were retained with an average of 2,319 unique molecular identifiers (UMI) and 1,419 genes per nucleus (**Extended Figure 1a**). Nuclei were integrated and clustered, accounting for batch effects (**Extended Figure 1b-d**). Cell type identification was performed using celltypist^33^, a semi-automated annotation pipeline followed by manual crosscheck with a curated list of gene markers^34–36^ (**Figure 1c**). To get a more fine-grained annotation, we performed sub-clustering of the endothelial cells (ECs), the myeloid and the fibroblast population. We identified five endothelial cell subpopulations—arterial, venous, lymphatic, endocardial, and capillary—each defined by distinct marker expression (**Extended Figure 1e**). The myeloid population was subdivided into two subgroups: Ccr2^−^/Lyve1^+^ tissue-resident MP and Ccr2^+^Lyve1^−^bone marrow-derived MP^37^ (**Extended Figure 1f**). In the fibroblast population we annotated activated fibroblasts based on the expression of *Postn and Meox1* (**Extended Figure 1g**).

The spatial transcriptomic dataset was obtained from the hearts of 12-week and 18-month-old mice and included five replicates per condition. To maximize the retention of spatial information, we performed a permissive quality control, resulting in a total of 15,838 spots with an average of 38,156 UMI and 4,350 genes per spot (**Extended Figure 2a-c**). To explore the spatial organization of the heart, we integrated and clustered the spots, accounting for both spatial information and batch effects (**Extended Figure 2d-f**). We identified 11 niches (**Figure 1b and Extended Figure 2f**), representing shared domains across sections. Principal component analysis of pseudo bulk counts showed that these niches were the second most significant source of variation (**Extended Figure 2g**).

**Figure 2.**
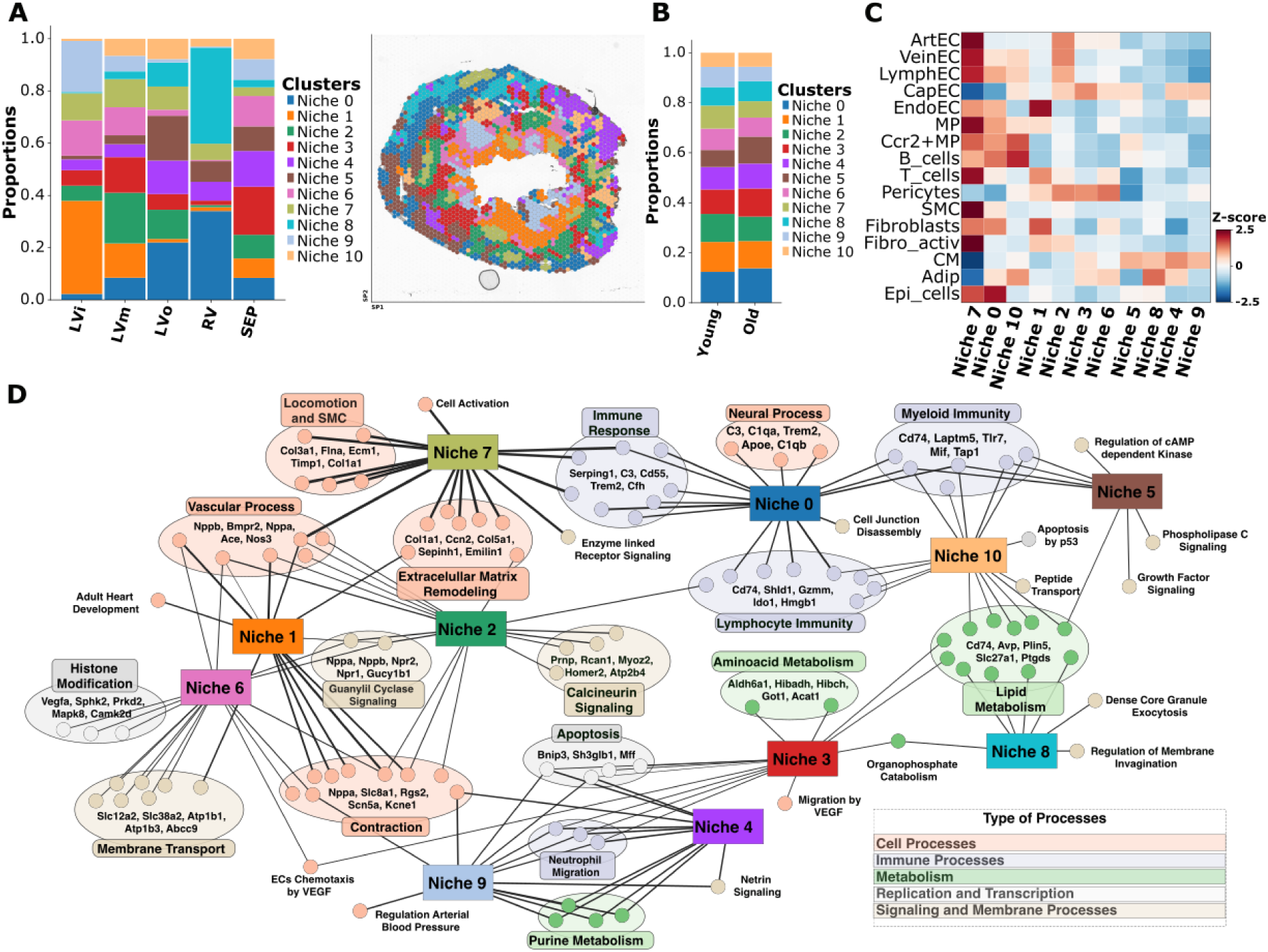
Characterization of cardiac niches. A) and B) Proportion of cardiac niches across A) anatomic regions and B) conditions. A) also shows the spatial distribution of the cardiac niches in a representative sample. Inner left ventricle (LVi); middle left ventricle (LVm); outer left ventricle (LVo); right ventricle (RV) and septum (SEP). C) Mean cell type proportion within each niche after Z-score transformation. The niches were hierarchically clustered. D) Network of the gene ontology terms enriched in each niche. The related terms were grouped and the top five genes shared with the highest score are shown.

To improve the resolution of the spatial transcriptomics, we integrated the data with the snRNA-seq using cell2location^20^; a deconvolution method that maps cell types and predicts their abundance in individual spots. Based on the predicted abundances, we calculated the proportion of each cell type per spot, observing expected distributions (**Figure 1d)**, with an average of 5 cells per spot. Cardiomyocytes (CM) were the most abundant cell type, accounting for 65 % of the total area (**Extended Figure 2h**), followed by capillary endothelial cells (CapEC) and pericytes. The pericyte to CapEC ratio was 1:2.4, reflecting previous histological findings^38^.

### Impact of aging on the cellular and molecular landscape of the cardiac niches

Aging can lead to a progressive deterioration in the structure and function of the heart; however, these changes may differ across regions. Therefore, we explored the structural, molecular, and cellular age-dependent alterations across the different cardiac niches. Their spatial distribution was examined by manually annotating three major anatomic regions: the right ventricle (RV), the septum (SEP), and the left ventricle. The left ventricle was subdivided into three sections: the inner (LVi), middle (LVm), and outer (LVo) ventricle (**Extended Figure 3a**). The left ventricle displayed a higher diversity in niche composition compared to the right ventricle (**Figure 2a**), reflecting the structural, physiological, and compositional differences of these regions^39^. Niches 1 and 9 were enriched in the endocardial region (**Figure 2a**). Niches 0 and 8 were predominantly found in the right ventricle. However, niche 0 showed an increased abundance towards the epicardial region of the left ventricle, a pattern also observed for niche 5. Niche 10 also showed an increase in abundance from the endocardial to the epicardial region of the left ventricle. The remaining niches were evenly distributed across all sections. With aging, the overall abundance of the cardiac niches remained comparable, except for niches 5, which showed an increase and 7, which showed a decrease in the aging heart. However, these changes were not statistically significant indicating that there was not a preference for specific regions during sample preparation (**Figure 2b and Extended Figure 3b, c**).

**Figure 3.**
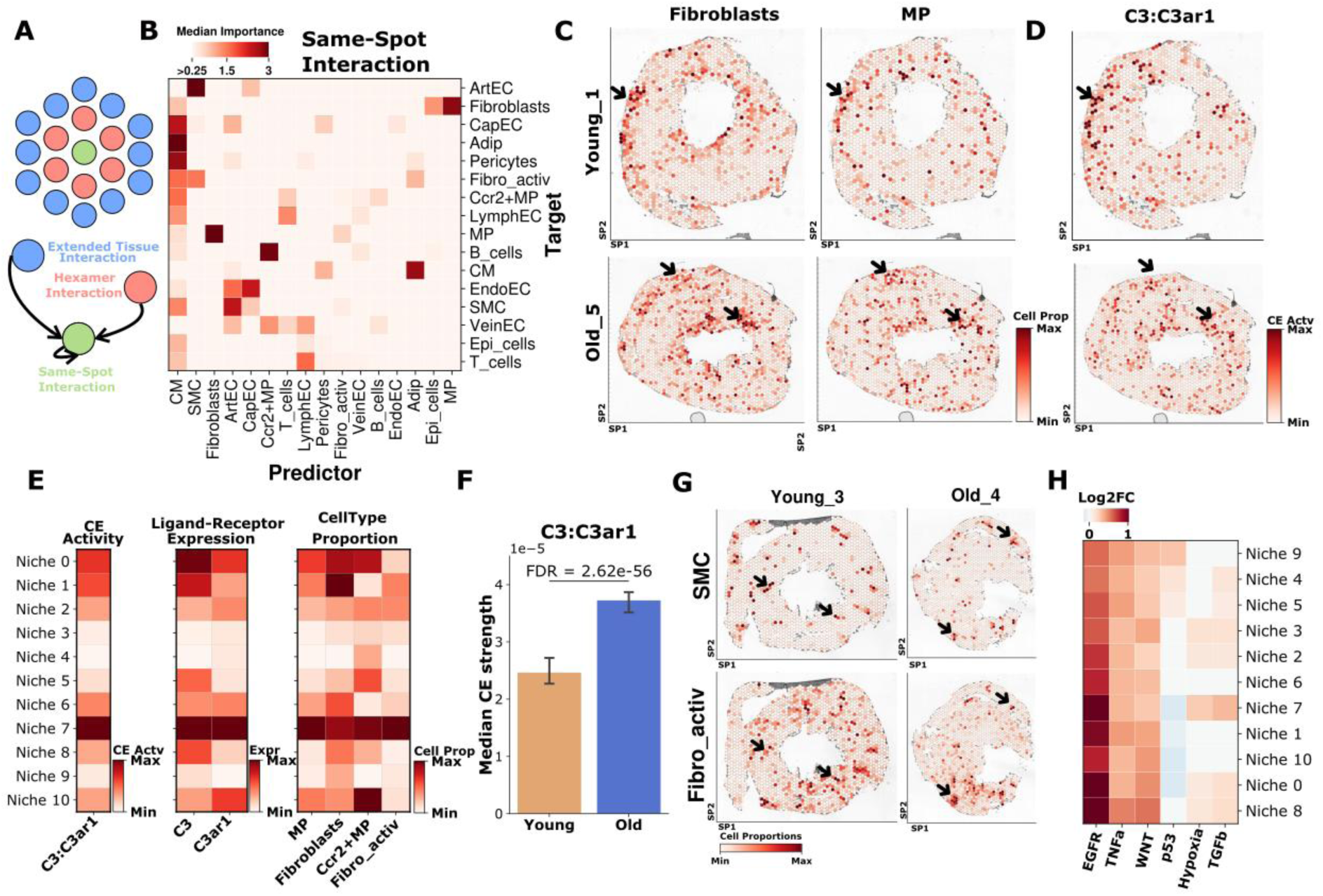
Cell interactions in the heart. A) Schematic on the type of interactions studied. B) Median importance over all samples of the cell types to predict the presence or absence of other cell types in the same spot. C) and D) Spatial distribution of C) the fibroblasts and macrophage (MP) proportion and D) the communication event (CE) activity of the C3:C3ar1 pathway for a representative young sample (top panel) and old sample (bottom panel). The arrows mark regions with overlap between the cell types and CE activities. E) Average CE activity (left panel), expression (middle panel) and cell type proportion (right panel) across all cardiac niches after scaling by subtracting the minimum and dividing by the maximum. F) Median CE for C3:C3ar1 in the spatial transcriptomics (Wilcoxon test p-value after Benjamini-Hochberg correction). G) Spatial distribution of smooth muscle cells (SMC) and activated fibroblasts in a representative young sample (left panel) and old sample (right panel). H) Log2 fold change (Log2FC) of pathways that showed an overall activation in the aged condition across niches.

#### Aging augments fibrotic gene expression and modulates macrophage populations in vascular niches

Next, we assessed the cellular composition of the niches and analyzed their functional differences using the predicted cell-type proportions and performing differential gene expression (DGE) analysis (**Figure 2c, d**). Arterial, venous, and lymphatic ECs were enriched in niche 7, as well as smooth muscle cells (SMC) and various inflammatory cell types. Consistent with these observations, processes related to vascular development and blood vessel morphogenesis were enriched (**Figure 2d and Supplementary Table 1**). This suggests that niche 7 likely represents ‘large vessel niches’, which are comprising arteries, veins, and lymphatic vessels. In the aged condition, an increase in collagen metabolism was observed **(Extended Figure 4a and Supplementary Table 2**), likely reflecting the perivascular fibrosis associated with aging^40^. Additionally, a metabolic switch was noted, transitioning from lipid catabolism and glycogen biosynthesis to glycolysis and lipid formation (**Supplementary Table 2**). Moreover, terms related to synapse pruning, microglia activation and proliferation, and neuron apoptotic process were elevated in the aged condition (**Extended Figure 4a**), aligning with the recently described neuro-repulsive environment of the aged vasculature^41^.

**Figure 4.**
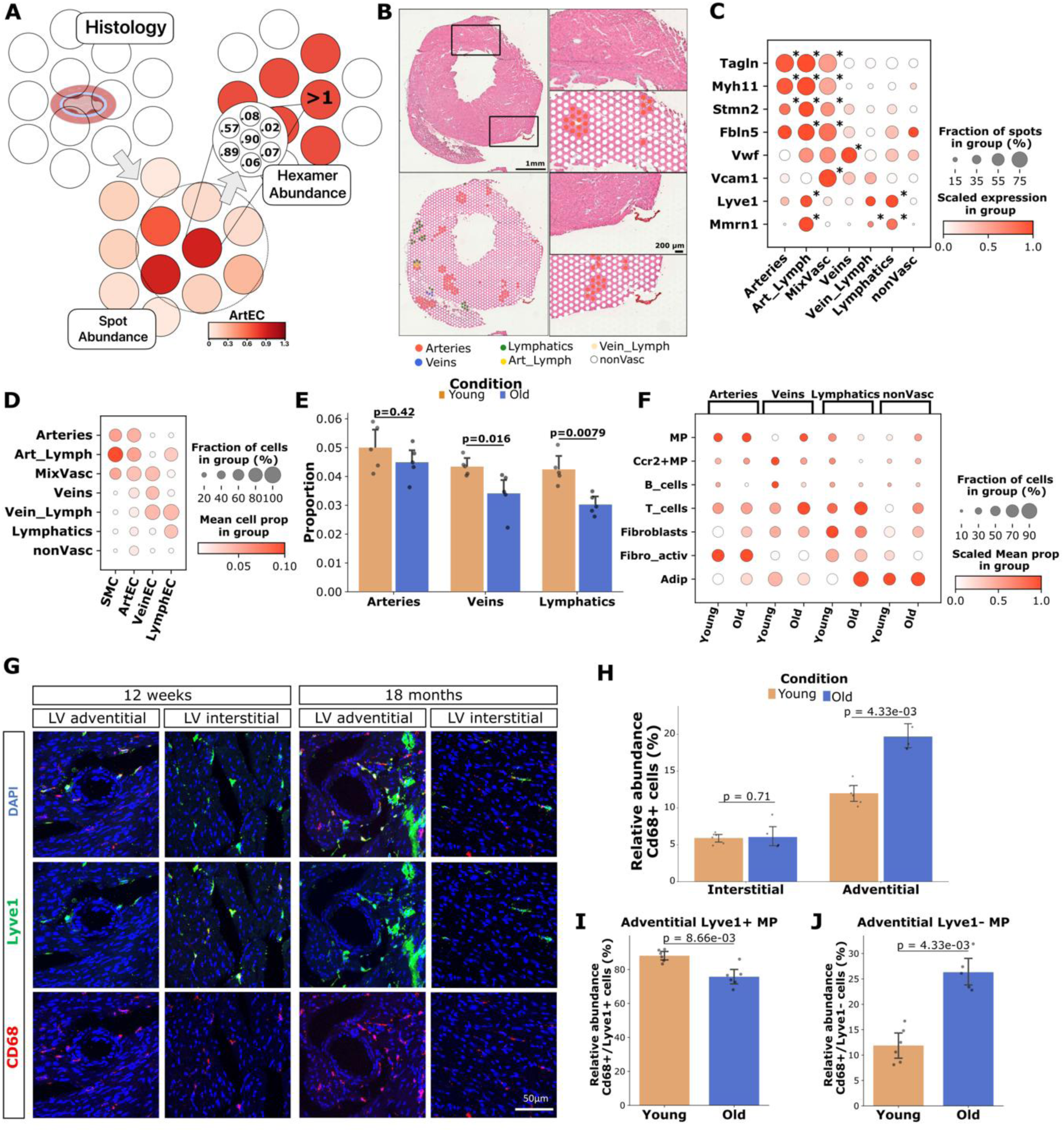
Compositional changes in the aging vasculature. A) Schematic on the approach employed to identify and annotate vessels on spatial transcriptomics. B) Haematoxylin and eosin staining (top panel) and vasculature annotation (bottom panel) in a representative slide. non-Vasc (non-vasculature). C) Average expression of marker genes across the annotated vessels after scaling to [0, 1]. The asterisk (*) represents the groups where the mean expression was significantly up-regulated compared to the non-vasculature spots (Wilcoxon test and Benjamini-Hochberg correction with adjusted p-value < 0.05 and Log2FC > 1). D) Mean cell proportion across the annotated vessels. E) Mean proportion per sample of artery, vein, and lymphatic endothelial cells in its representative vessels. The error bar represents the 95 % confidence interval of the mean. Significance was tested with the Wilcoxon Rank Sum test. F) Average cell proportion in each type of vessel for each condition scaling like in C). To calculate the fraction of cells in the group, we excluded spots with a proportion < 0.01. G) Immunofluorescence staining of a representative old and young male mice sample from the adventitial and interstitial region. CD68 (red, macrophages); Lyve1 (green, resident macrophages and lymphatics) and DAPI (dark blue, nucleus). Scale bars, 50 µm. H) Relative abundance of Cd68^+^ cells in interstitial and adventitial region for young and old mice samples. I) and J) Relative abundance of I) Lyve1^+^ and Cd68^+^ cells and J) Lyve1^−^and Cd68^+^ cells. In H), I) and J) the error bars represent the 95 % confidence interval of the mean. Significance was tested with the Wilcoxon Rank sum test and the error bar represents the 95 % confidence interval of the mean.

The cellular composition of niche 0 was comparable to niche 7; however, instead of arterial ECs, it was predominantly enriched in epicardial cells, reflecting its spatial enrichment in the epicardial region of the heart (‘Epicardial niche’). This niche was characterized by immune-related processes, particularly antigen presentation, which were pronounced in the aged condition (**Extended Figure 4b**). In contrast, processes related to endothelial cell remodeling and epithelial cell development were enriched in the young condition (**Extended Figure 4b**).

Consistent with its spatial localization to the endocardial region, niche 1 was enriched in endocardial ECs, and was identified as the ‘endocardial niche’. This niche was characterized by an enrichment of genes related to the regulation of heart rate and blood pressure. In the young hearts, genes associated with junctions, focal adhesion, and the establishment of the endothelial barrier were prominent (**Extended Figure 4c**). In the aged heart, processes related to microglia activation and macrophage polarization stood out.

Niche 2 comprises primarily larger vessel ECs, with a lower presence of SMC and a higher enrichment of pericytes (**Figure 2c**), resembling regions that are a mixture of small arterioles and venules. This niche was characterized by processes related to muscle cell development and adaptation, with an overrepresentation of vascular remodeling processes in the young condition. In contrast, in the aged condition, it was marked by genes associated with microglia activation (**Extended Figure 4d**).

Niche 10, which was particularly enriched in the septum and left ventricle, comprises immune cells, adipocytes, and, to some extent vascular ECs (**Figure 2c**). In line with this observation, the niche showed enrichment in processes related to myeloid immunity and fatty acid metabolism. Notably, it also displayed an overrepresentation of processes related to microglia cell activation and macrophages in the aged condition (**Extended Figure 4e**). Terms related to collagen metabolism were also observed.

In general, all vessel-associated vascular niches (0, 1, 2, 7, and 10) exhibit similar gene enrichment patterns in the old heart, with a surprising inclusion of gene ontology terms associated with microglial cells. Although the heart does not contain microglia, it does harbor a type of immune cell, the resident and bone-marrow-derived MP, which may share functional similarities. We observed an enrichment of Lyve1+ MP and bone marrow-derived Ccr2+MP in these niches, particularly in the aged condition (**Figure 2c and Extended Figure 4f**). In addition, *Grn*, *Trem2,* and *Tyrobp*, associated with the microglia terms, were also predominantly expressed in both resident and bone marrow-derived MP (**Extended Figure 5a**). DGE analysis in the Ccr2+MP in the aged hearts revealed upregulation of genes such as *Fgr* and *Myof1* (**Supplementary Table 3**), which have previously been linked to a pro-inflammatory phenotype^42,43^. Tissue-resident MP also showed enrichment for genes that have been linked to a pro-fibrotic phenotype, including *Cd209*, *Timp2*, and *Ccl8*^44,45^ (**Supplementary Table 3**). These genes were also enriched in vascular niches (**Extended Figure 5b**). Overall, this suggests a complex microenvironment in the vascular niches with macrophages contributing both to a pro-inflammatory and pro-fibrotic environment.

**Figure 5.**
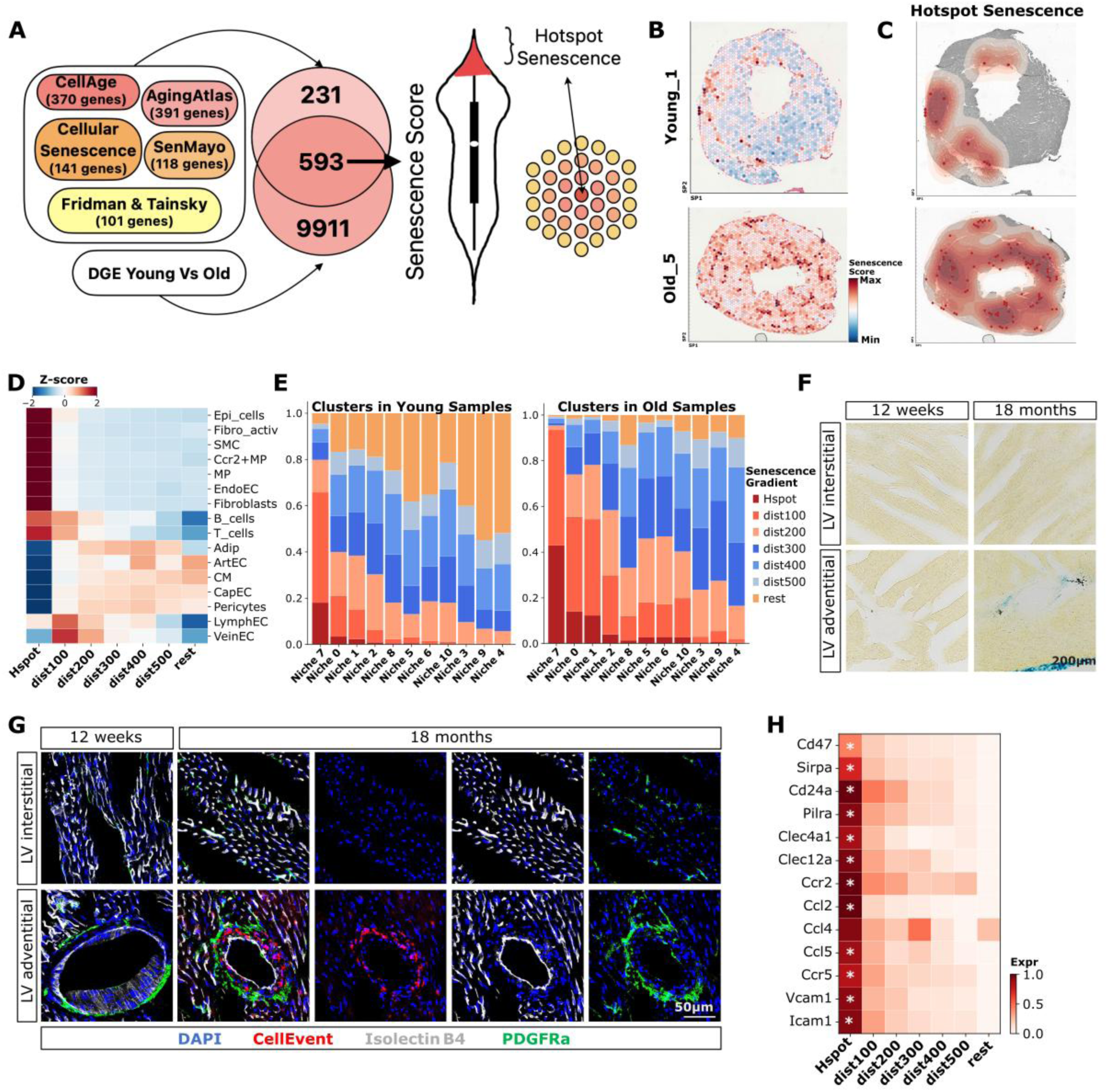
Cellular senescence in the aging heart. A) Schematic on how the hotspots of senescence (Hspot) have been defined. B) and C) Spatial distribution of B) the senescence score and C) the density of the Hspots in two representative samples. D) Median cell proportion after Z-score transformation in the Hspots and spots located at different distances (in µm). E) Proportion of Hspots and spots located at different distances (in µm) across clusters in young and old samples. F) and G) SA-ß-galactosidase staining and immunofluorescence showing senescence in a representative interstitial and adventitial region in young and old male mice hearts. Isolectin B4 (white, endothelial cells); PDGFRa (green, fibroblasts); CellEvent (red, senescence) and DAPI (dark blue, nucleus). Scale bars, 200 µm in F) and 50 µm in G). H) Average expression of genes associated to the immune inhibitory pathway and recruitment of immune cells after standardize scaling to [0, 1]. The asterisk (*) represents the groups where the mean expression was significantly up-regulated compared to the non-vasculature spots (Wilcoxon test and Benjamini-Hochberg correction with adjusted p-value < 0.05 and Log2FC > 0.25).

#### Capillary and cardiomyocyte niches undergo remodeling and alterations of the circadian clock genes

Niche 3, consisting of capillary ECs and pericytes was primarily localized to the septum and the middle region of the left ventricle. Notably, cardiomyocytes were also enriched here. This niche displayed an enrichment for processes related to catabolism, suggesting a higher metabolic activity. In the aged heart, it was characterized by genes associated with hepatic stellate cell activation, including *Acta2*, *Cygb,* and *Rps6ka1* (**Extended Figure 5c and Supplementary Table 2**). Since activation of hepatic stellate cells occurs during liver fibrosis, this may reflect fibrotic pericyte activation, including the secretion of extracellular matrix proteins. Niche 6 had a similar cellular composition as niche 3 but showed lower cardiomyocytes abundance and a higher presence of other cell types, such as fibroblasts. In the aged hearts, this niche exhibited increased membrane trafficking activity and processes related to cardiac muscle repolarization (**Extended Figure 5d**). Both capillary enriched niches showed enrichment for processes related to extracellular matrix remodeling in the young condition with alterations in the cell-cell junctions.

Niches 4, 5, 8, and 9 were enriched in cardiomyocytes, with niches 4 and 9 showing a higher presence of capillary ECs and cardiomyocytes but not pericytes (**Figure 2c**). This suggests that these niches represent ECs in proximity to cardiomyocytes. Interestingly, these niches can be mapped to distinct spatial locations: niche 4 was localized to the epicardial region and the septum, while niche 9 was primarily found in the endocardial region (**Figure 2a**). Niche 8 was mainly localized to the right ventricle, and niche 5 was found in the epicardial region (**Figure 2a**). Niches 4, 8, and 9 were characterized by an enrichment of metabolic processes (**Figure 2d**). We observed that the capillary and cardiomyocyte-enriched niches showed an overrepresentation of processes associated with cardiac muscle repolarization and circadian clock (**Extended Figure 5e-h and Supplementary Table 2**). In contrast, the young hearts were enriched in processes associated with remodeling, such as regulation of tight junction assembly, connective tissue replacement, and cell development (**Extended Figure 5e-h**).

### Cellular interactions and macrophage activation in the vascular niche of the aging heart

Our analysis of the cardiac niches revealed an enrichment of a pro-fibrotic macrophage phenotype particularly in the large vessel niche. To further understand the associations between cell types and potential interactions leading to macrophage activation, we employed mistyR^25^, a framework designed to assess the relationship between cell types by estimating their importance in explaining the presence or absence of other cell types. The estimated importance can reflect co-localization or co-exclusion of two cell types. To gain further insights into changes by distances, we analyzed three cases: 1) same-spot interactions, 2) hexamer space (direct neighbors) interactions, and 3) extended tissue space interactions (**Figure 3a, b and Extended Figure 6a, b**).

As expected, we observed that the absence of cardiomyocytes was associated with an increased presence of other cell types, highlighting the central role of stromal cells in tissue organization (**Figure 3b and Extended Figure 6c**). In line with the known anatomy of the vasculature, SMC were particularly associated with arterial ECs, while pericytes coincided with capillary ECs, even when extending the analysis to the hexamer space (**Extended Figure 6a, b**). We found differential interaction among inflammatory cells: tissue-resident MP were associated with fibroblasts, showing a co-localization that was more pronounced in the aged condition (**Figure 3c**). Bone marrow-derived MP showed co-localization with B cells (**Extended Figure 6d**). Inference of cell communication events using HoloNet^26^, predicted activation of the C3:C3ar1 pathway in regions enriched in tissue resident MP and fibroblasts (**Figure 3d, e**), with a significant upregulation in the aged heart (**Figure 3f**). The snRNA-seq further supported a possible interaction of both cell types, since fibroblasts expressed the ligand (*C3*), and MP expressed the receptor (*C3ar1*), both of which were significantly upregulated in the aged condition (**Extended Figure 6e**). Lastly, we also observed co-localization between activated fibroblasts and SMC (**Figure 3b, g**). However, the influence of SMC on the distribution of activated fibroblasts decreased with increasing distance. Conversely, MP distribution emerged as a more significant predictor (**Extended Figure 6a, b**).

### Pathway activity dynamics across the cardiac niches

We next investigated pathway activities using PROGENy^21^. Our analysis revealed a notably stronger activation of the EGFR pathway in the aged cardiac niches (**Figure 3h**), particularly within the epicardial region and the vasculature (niches 8, 0, and 7). Additionally, the WNT and TNF*α* pathways were enriched in the aged heart. Hypoxia-related gene signatures were upregulated in niches enriched with capillary ECs (niches 2 and 3), as well as in niche 7. TGF-β signaling also showed higher activation in the aged heart, particularly in niche 7, which aligns with our cell interaction analysis that indicated co-localization between SMC and activated fibroblasts. Finally, the tumor suppressor gene p53, known for its role in maintaining cardiac homeostasis^46^, was downregulated in most aged niches, except for niches 4, 5, and 9, which were enriched in cardiomyocytes.

### Identification and annotation of large vessels

Our characterization of the cardiac niches identified niche 7 as a vascular niche comprising all the major large vessels of the heart. However, this niche was not the only one associated with vasculature and this analysis did also not allow us to distinguish among the different subtypes of vessels. In mice, arteries range from 150-160 µm in diameter, while veins can reach up to 250 µm^47,48^. The per-spot capture area of the spatial transcriptomic platform was 55 µm in diameter with 100 µm between two spots. This can lead to the partial capture of large vessels. To address this limitation, we extended the analysis to direct neighbors in the hexamer space (**Figure 4a**). We aggregated the estimated cell abundance for the different cell types considering a maximum radius of 100 µm, which represent direct neighboring spots. Next, we binarized the matrix, where a cell type is considered present if the aggregated abundance exceeds one, effectively interpreting this as the presence of at least one cell. The binarized matrix is then used to annotate spots containing the large vessels. For each type of vessel, at least one EC subtype was detected within the hexamer space. Additionally, for arteries, we imposed another constraint, requiring the presence of at least one SMC.

After applying our strategy, we were able to differentiate arteries, veins and lymphatic vessels (**Figure 4b and Extended Figure 7a**). However, due to the limited resolution, vessels in proximity resulted in a mixed cell compositions and annotation in some clusters (**Figure 4c, d**). The mixed clusters contained arteries and veins (MixVasc), arteries and lymphatics (Art_Lymph), and veins and lymphatics (Vein_Lymph). These clusters were excluded from subsequent analysis. DGE analysis between each type of vessel and the non-vascular spots revealed an enrichment of marker genes characteristic of these regions (**Figure 4c**). SMC and artery ECs markers (*Tagln*, *My11*, *Stmn2,* and *Fbln5*) were enriched in arteries; vein ECs markers (*Vwf* and *Vcam1*) in veins, and lymphatic ECs markers (*Lyve1* and *Mmrn1*) in lymphatics. Overall, our strategy provides an accurate supervised approach for identifying and annotating large anatomical structures accounting for differential cell type composition.

### Age-related changes in vascular composition and cell dynamics

Next, we examined how aging affects the different types of vessels, excluding spots with mixed cellular composition. Quantification of the vessel types revealed a tendency for reduced vascularization, with a significant decrease observed for veins and lymphatics (**Figure 4e)**. To understand the compositional changes in these regions, we assessed the overrepresented cell types (**Figure 4f**). Tissue-resident and bone-marrow derived MP were notably enriched in the vascular regions. Therefore, we performed immunostaining to quantify MP in the adventitial and interstitial region (**Figure 4g**). We observed an enrichment of Cd68+ cells, which was significant (p-value = 6.5e-12, ANOVA test), in the adventitial region compared to the interstitial region (**Figure 4h**). We did not observe a significance difference in the MP composition in the interstitial region between conditions, however, in the adventitial region we observed a switch upon aging from Cd68+/Lyve1+ cells representing resident MP and Cd68+/Lyve1-cells representing bone-marrow derived MP (**Figure 4i, j**). Aged lymphatic vessels exhibit a lower abundance of MP, which could indicate a decline in lymphatic drainage function during aging.

It has been reported that myocardial B cells typically remain in the intravascular region, with only a few crossings through the endothelium^49^. In our study, we observed an enrichment of B cells in the vascular region of the young heart, particularly in the veins (**Figure 4f**). T cells and adipocytes were present across the different regions, with a noticeable increase in abundance in the aged condition (**Figure 4f**). Lastly, fibroblasts were particularly enriched in the vascular region, with arteries showing a higher number of activated fibroblasts (**Figure 4f**).

### Senescence is predominantly induced in the vascular niches

Cellular senescence, a hallmark of aging, constitutes a stress response that is needed during normal development and tissue homeostasis. However, aging triggers mechanisms associated with an increase in cellular senescence leading to functional and structural changes in the microenvironment of senescent cells. To explore the spatial distribution of senescence in the aging heart, we identified hotspots of senescence (**Figure 5a**). Using established senescence gene sets^22,23,29–32,50^ and differentially expressed genes between young and aged samples, we compiled a list of 593 aging-sensitive genes (**Supplementary Table 4**). We then scored the spots based on the expression of these genes (**Figure 5b, c and Extended Figure 7b**) defining hotspots of senescence (Hspot)^51^. As previously reported^51^, the senescence score was highest in these hotspots and decreased with distance (**Extended Figure 7c**). We noted that while senescence in young samples was detected in some regions, in aged hearts the average score was higher and extended to a larger area (**Extended Figure 7c**). Analysis of cell compositions revealed that the senescence hotspots were particularly enriched in fibroblasts, SMC, and MP (**Figure 5d**). With aging, the cellular composition of these hotspots broadens: aged samples showed an additional increase in the proportion of cardiomyocytes and capillary ECs, reflecting a spreading of the senescence (**Extended Figure 5d, e**). Further analysis of the niches (**Figure 5e**) confirmed that the senescence hotspots were primarily localized to the vascular niche (niche 7). In aged samples, senescence extended into the endocardial niche (niche 1) and the epicardial region (niche 0). Examining the distribution in different vessel subtypes, we found that arteries were the primary source of senescence, followed by veins and lymphatics (**Extended Figure 7f**). Additionally, we observed a higher abundance of senescence hotspots in niches enriched in immune cells (niches 0 and 1). To validate these findings, we evaluated cellular senescence by performing SA-β-galactosidase and immunofluorescence stainings (**Figure 5f, g and Extended Figure 7g**). Consistent with the results obtained by spatial transcriptomics, we observed an increased localization of senescence cells in the adventitial of large vessels (**Figure 5f, g**). Interestingly, we identified a correlation between the Hspots and genes associated to the recruitment of immune cells, and genes associated to immune inhibitory pathways, including the genes *Cd47*, *Sirpa*, *Cd24a* and *Clec4a* **(Figure 5h**). Specifically, we observed an increase in the proportion of cells expressing genes associated to immune inhibitory pathway in the aged condition (**Extended Figure 7h**).

## Discussion

Aging is the result of the accumulation of molecular and cellular damage over time, which in turn results in structural and molecular alterations, as well as changes in the cellular compositions of organs. Integration of spatial transcriptomics data across young and aged hearts allowed us to identify 11 cardiac niches, which were characterized by distinct cellular composition and functional signature. Furthermore, the integration with snRNA-seq allowed us to study the changes in the cellular dynamics. Aging did not change the cardiac niches proportions. However, we observed distinct regional changes across the different regions; with the left ventricle showing a higher diversity and exclusive enrichment for some niches like niche 9 and 6, which may reflect the different developmental origin^52^ and distinct pressure levels between the ventricles.

Endothelial cells, which are the most abundant non-myocyte cell type in the heart^53^, form a network that is vital for tissue perfusion, ensuring proper oxygenation and nutrient delivery to the myocardium. One key novel finding of this study is that age-associated changes primarily impacted the vascular niches, particularly the niches associated to large vessels, which exhibited both pro-fibrotic and senescence signatures. This pro-fibrotic environment is reflected on the upregulation of collagen metabolism, enrichment of activated fibroblasts, and activation of TGF-β signaling. In addition, both resident and bone-marrow derived MP, showed co-localization with fibroblasts within the vascular niches. We also observed a switch in the MP population around the vessels, with a decrease of resident MP and an increase of bone-marrow derived MP; suggesting a complex inflammatory environment that may contribute to age-associated vascular dysfunction. Among the cellular communication pathways which are enriched in aging, the complement C3:C3ar1 interaction stood out. Complement *C3* and its receptor *C3ar1* play complex roles in cardiac function, exhibiting both protective and potentially detrimental effects depending on the context and site of activation. While *C3* provides protective effects in ischemia/reperfusion injury models^54^, it contributes to right ventricular dysfunction and fibrosis^55^. Our study predicts that fibroblast-derived *C3* activates *C3ar1* in MP within the aging heart. Notably, *C3ar1* promotes MP polarization toward an M2 (anti-inflammatory) phenotype^56^. While this may help suppress inflammation, M2 macrophage are also known to drive fibrosis, a hallmark of age-related pathology ^57^. Recent studies suggest that cardiac fibroblasts exhibit functions in innate and adaptive immunity in acute injury mouse models and can aggravate the inflammatory response^58^. It is interesting to note that we detect similar activation pattern in the aging heart, specifically in the vascular niches, but less so in the interstitial regions. This indicates distinct roles and activation patterns between perivascular and interstitial fibroblasts in the myocardium.

Besides the pro-fibrotic and pro-inflammatory environment, vascular niches were identified as hotspots of senescence. This is also supported by the metabolic shift towards glycolysis and lipid formation observed in these regions, which is consistent with senescence-associated metabolic alterations^59,60^. Therefore, one might envision a vicious cycle which supports progression of an environment around the vasculature, which promotes inflamm-aging and accumulation of senescence.

Under normal circumstances, senescent cells are removed by MP and other immune cells through a process called immunosurveillance^61^. However, clearance of the senescent cells by MP can be disturbed by the expression of immune inhibitory molecules, which allow cells to elude MP phagocytosis and immune surveillance^62^. We observed ‘immune evasion’ genes, including *Cd47*, *Sirpa*, *Cd24a*, and *Clec4a*^63–65^, enriched in the hotspots of senescence and their expression declined related to their distance from the hotspots of senescence.

Surprisingly, we also identified senescence hotspots in young hearts that were not recapitulated by histological analysis. This discrepancy could be due to several factors, including the arbitrary selection of the 5% threshold for defining senescence and the specific genes used in our senescence score. Another possibility is that cells with transcriptional activation of senescence genes are rapidly cleared in young hearts and are therefore no longer detectable by immunostaining. Interestingly, however, the few senescence hotspots identified through *in silico* analysis in young hearts are also located near blood vessels, suggesting that these regions may have an increased susceptibility to acquiring a senescent state.

We also observed age-associated changes in cardiomyocyte-enriched niches, showing a disruption of circadian clock genes. Circadian genes are critical for cardiovascular physiology, and aging has been shown to disrupt the circadian patterns^66^. The dysregulation of clock genes in the cardiac niches may contribute to age-related cardiac electrical disturbances and arrhythmias, which aligns also with the enrichment of muscle repolarization processes in these niches. Previous studies have shown that the disruption of clock function in murine cardiomyocytes is linked to alterations in electrical conduction, thereby increasing susceptibility to arrhythmia^67^.

Besides the stronger activation of TGF-β in vascular niches, the analysis of other signaling pathways showed stronger activation of EGFR, WNT, TNF*α*, while p53 showed less activation in some niches. The chronic activation of these pathways has been implicated in pathophysiological cardiac remodeling. For example, TNF*α* levels correlate with the severity of heart failure^68^ and can induce left ventricular dysfunction and cardiomyopathy^69^. Chronically high activation of the WNT/ β-catenin has a pro-fibrotic and hypertrophic effect in the adult heart^70^. On the other hand, downregulation of p53 has been linked to age-related cardiac hypertrophy and heart failure^71^. Altogether, the dysregulation of these pathways highlights regions that are susceptible for pathophysiological cardiac remodeling.

This study has several limitations. First, the spatial transcriptomics platform lacks true single-cell resolution, restricting our ability to precisely attribute observed molecular changes to specific cell types. Second, most of our findings are based on gene expression analysis rather than protein expression, which may not always align^58^ and does not account for the many post-transcriptional regulatory mechanisms. While we validated some findings through histological staining, the causal role of the dysregulated genes has not been assessed.

In summary, our findings reveal the intricate interplay of structural, cellular, and molecular changes across cardiac niches during aging. Interestingly, our results suggest that aging predominantly affects vascular niches, reinforcing the notion that the vasculature plays a central role in organ aging. This is further supported by evidence that systemic treatment with vascular endothelial growth factor promotes healthy aging and extends lifespan^72^. The enrichment of macrophages and activated fibroblasts and senescent cells in the age-associated vascular niche further underscores the critical role of immune-fibroblast interactions in cardiac aging. Additionally, the observed circadian disruptions highlight potential therapeutic targets for mitigating age-related cardiac dysfunction. Future studies should aim to dissect the mechanistic drivers of these niche-specific alterations and explore targeted interventions to preserve cardiac function with aging.

## Nonstandard abbreviations and acronyms

CapEC: capillary endothelial cells
CE: communication event
CM: cardiomyocytes
DGE: differential gene expression
EC: endothelial cells
LVi: left ventricle inner
LVm: left ventricle middle
LVo: left ventricle outer
MP: Macrophages
UMI: unique molecular identifiers
RV: right ventricle
scRNAseq: single-cell RNA-sequencing
snRNAseq: single-nucleus RNA-sequencing
SEP: septum
SMC: smooth muscle cells

## Acknowledgments

SD is supported by the ERC advanced grant Neuroheart and the German Research Foundation (DFG; SFB 1366 (Project number 394046768), Project B04). SD and MHS are supported by the SFB 1531 (Project number 456687919, Project B01 and S03). SD, LT, and CK are supported by the DZHK shared expertise project.

## Author contributions

D.R.M. and S.D. designed the research; D.R.M. and M.R.J. performed and supported bioinformatic analysis; M.H.S. and D.J. supervised the bioinformatic studies; V.L. performed histological analysis, L.T., L.Z., and C.K. performed and supported analysis of spatial transcriptomics; A.M.Z. contributed to the biological interpretation of the results. D.R.M., and S.D. wrote the paper. All authors read and approved the final version of the paper.

## Materials & Correspondence

Correspondence and material requests should be addressed to Prof. Stefanie Dimmeler.

## Novelty and significance

### What is known?

- Aging is associated with increased senescence and a decline in physiological processes, leading to an increased risk for developing cardiovascular diseases.
- Endothelial cells are central components in the heart forming a vital network for tissue perfusion, ensuring proper oxygenation and nutrient delivery to the myocardium.

### What new information does this article contribute?

- The manuscript provides spatial transcriptomics data from young and aged murine hearts.
- Aging affects primarily the perivascular regions.
- Aging is associated with an interaction of macrophages and fibroblasts through the C3:C3ar1 axis.
- Vascular niches are the primary hotspots of senescence with increased expression of immune inhibitory molecules.

We generate spatial transcriptomics from young and aged murine hearts and increased the resolution by integrating it with single-nucleus RNA sequencing. We map all the major cell types present in the heart and studied changes in their spatial distribution during aging. Additionally, we identified cardiac niches with distinct distribution and signatures. We identified the large vessel-associated vascular niches as the primary hotspot of age associated changes. The vascular niches were characterized by a pro-inflammatory and pro-fibrotic environment with increased abundance of bone-marrow derived macrophages and activated fibroblasts. Senescence was also particular enriched in these regions with a co-localization of immune inhibitory molecules promoting the immune cell evasion.

---

**Extended Figure 1.**
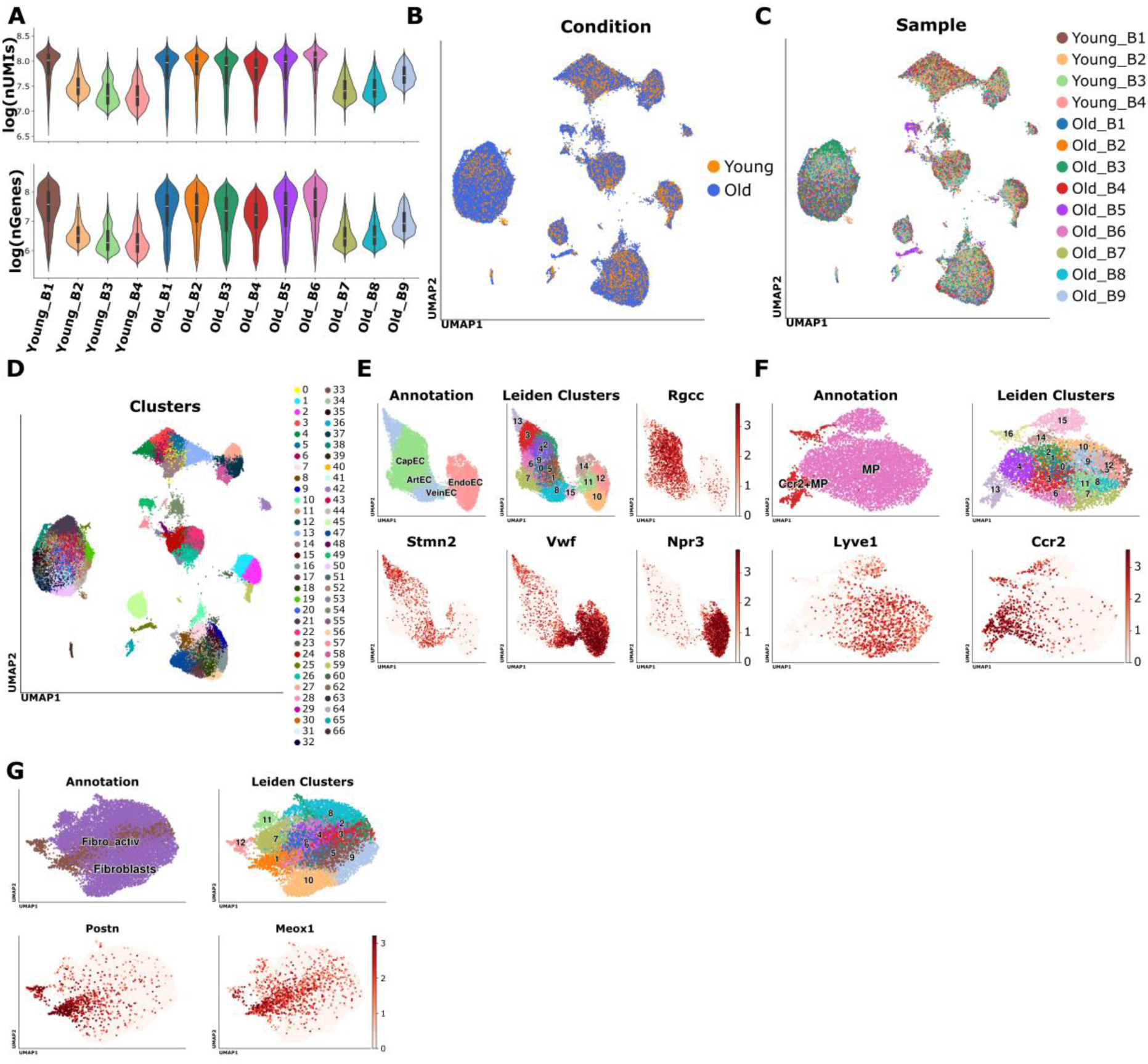
snRNA-seq quality measurements. A) Distribution of the total number of unique molecular identifiers (UMI) and number of genes per nucleus detected across samples in logarithmic scale. B), C) and D) UMAP of all the cells splitting by B) condition, C) sample and D) clusters identified by the leiden algorithm. E), F) and G) UMAP embedding of the sub-clustering showing the annotation, the clusters and the expression of marker genes for E) endothelial cells, F) myeloid cells and G) fibroblasts.

**Extended Figure 2.**
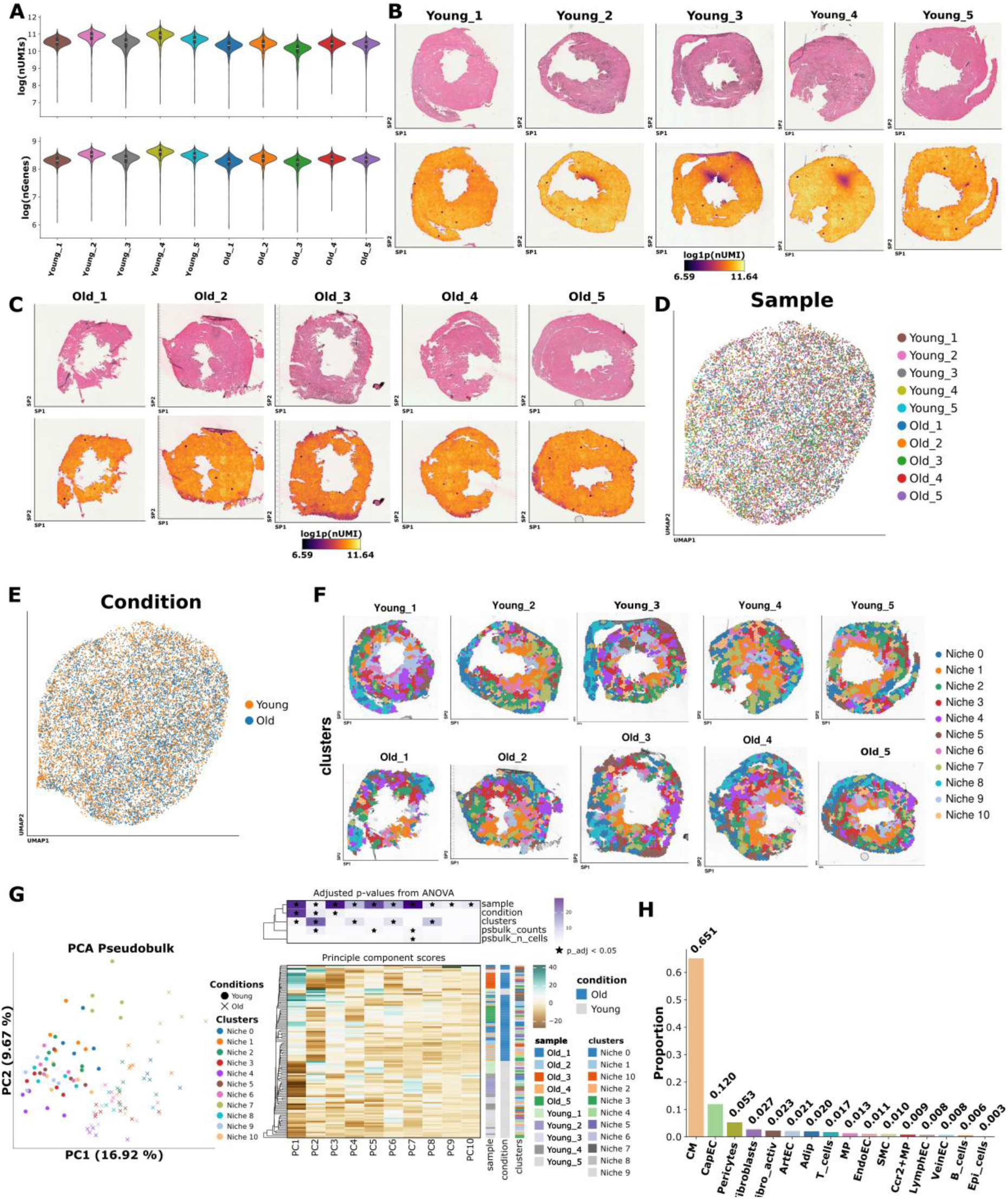
Spatial transcriptomics quality measurements. A) Distribution of total number of unique molecular identifiers (UMI) and number of genes per spot across samples in logarithmic scale. B) and C) Hematoxylin and eosin staining (upper panel) and spatial distribution of UMI counts in logarithmic scale (lower panel) for B) young samples and C) old samples. D) and E) UMAP of all the samples splitting by D) samples and E) condition. F) Spatial distribution of the clusters. G) Principal component (PC) analysis representation of the pseudobulk counts and heatmap showing the distribution of the scores across all the spots for each PC. The association between each PC and metadata was tested with ANOVA. P-values were corrected with the Benjamini-Hochberg method. H) Proportion of cell types across all samples.

**Extended Figure 3.**
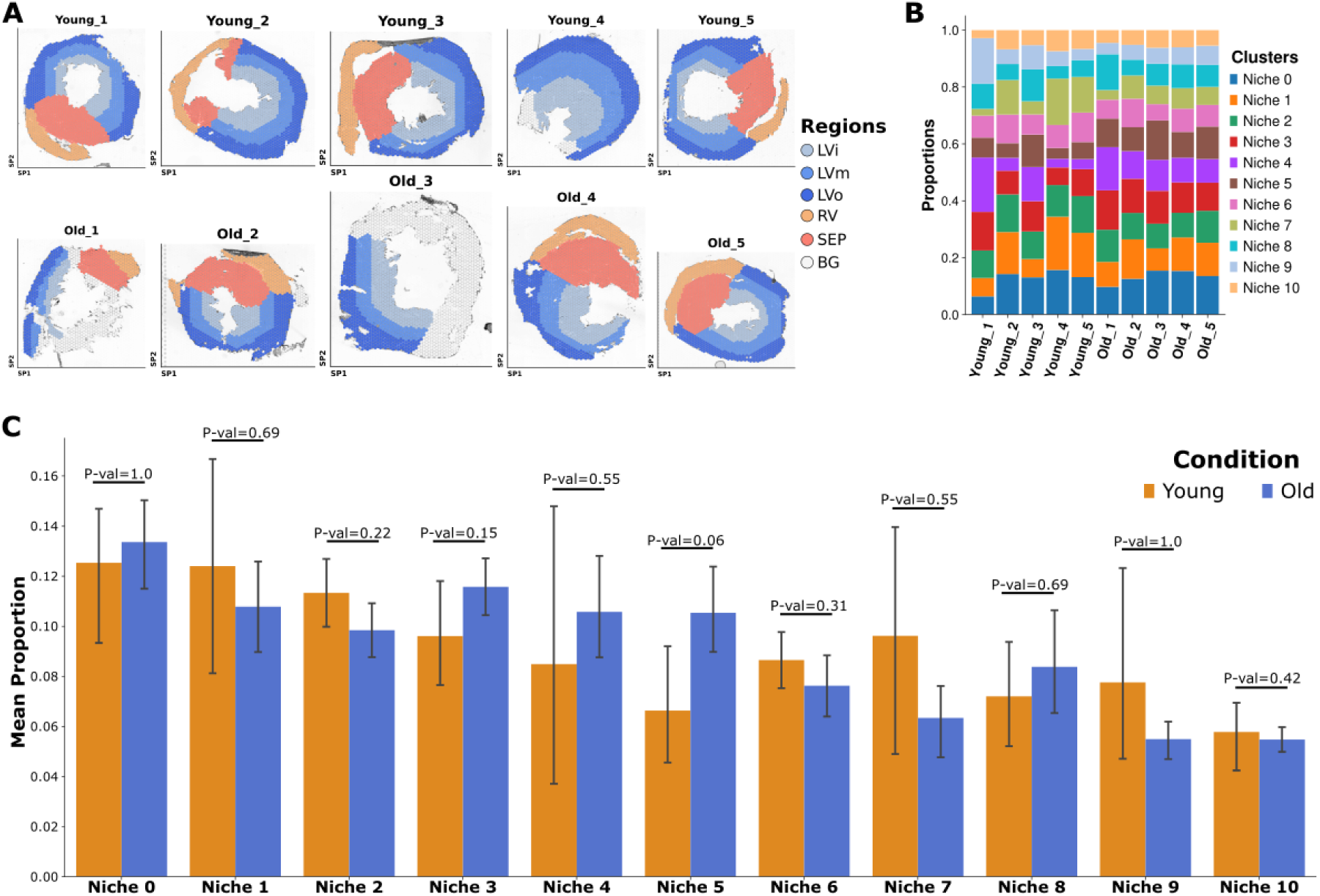
Structural and compositional analysis of the spatial transcriptomics. A) Illustration showing the spatial distribution of the manual annotation of the anatomic regions B) Proportion of each niche per sample C) Mean proportion of niches per condition. The p-values were estimated with Wilcoxon test comparing the sample proportions between both conditions. The error bar represents the 95 % confidence interval of the mean.

**Extended Figure 4.**
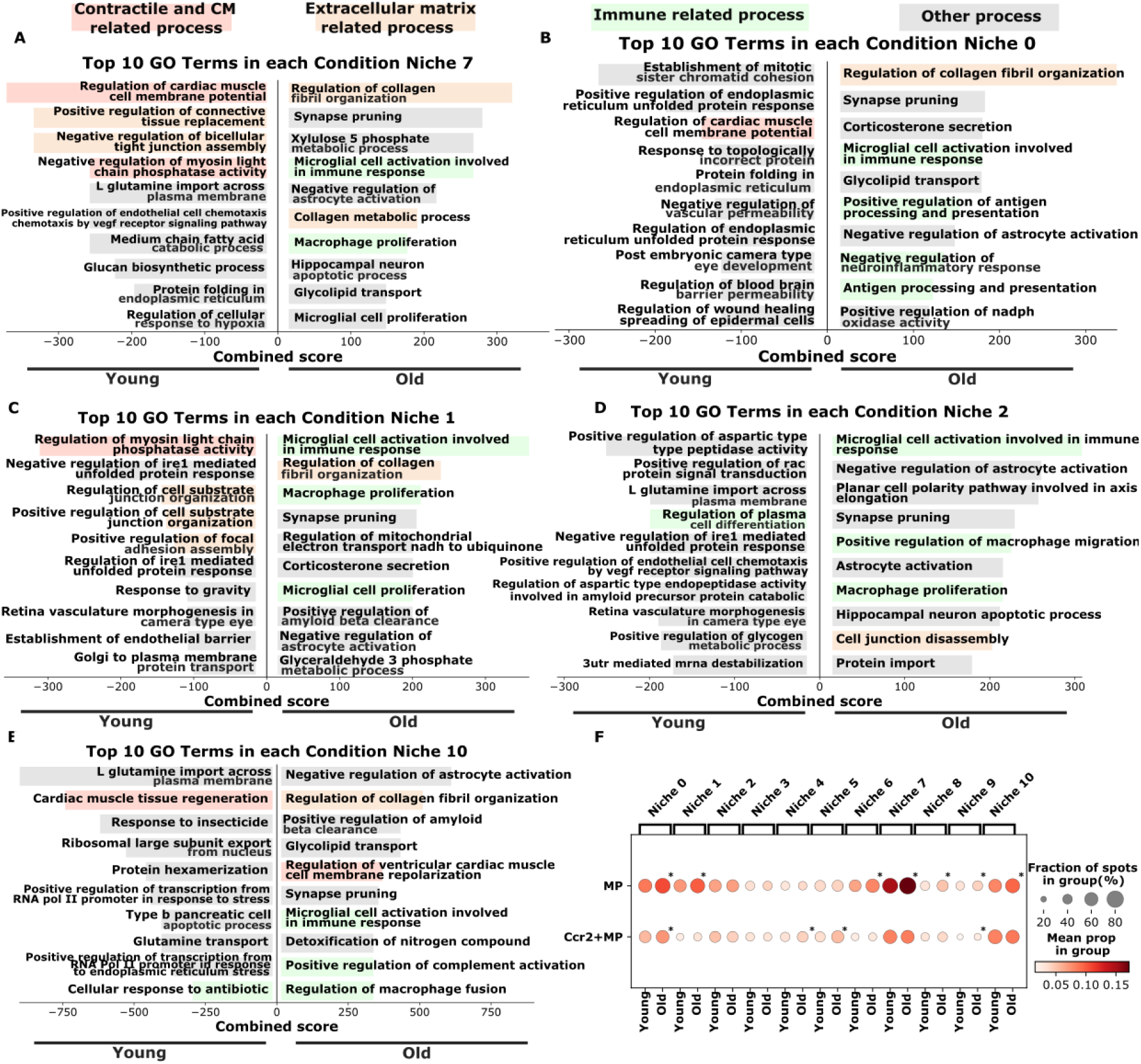
Functional analysis of the cardiac niches enriched in endothelial cells. A) to E) shows the top gene ontology terms regulated in the old (positive axis) and in the young condition (negative axis) for each cardiac niche (7, 0, 1, 2, and 10). The terms were filtered using an adjusted p-value of 0.05 and ranked base on the combined score estimated by enrichR. Terms associated with similar processes were grouped and colored together. F) Mean proportion of macrophages (MP) and Ccr2+MP. The asterisk (*) indicates that the proportion of cell types is significantly higher in the old condition within each niche (adjusted p-value < 0.05; using Wilcoxon test and Benjamini-Hochberg correction).

**Extended Figure 5.**
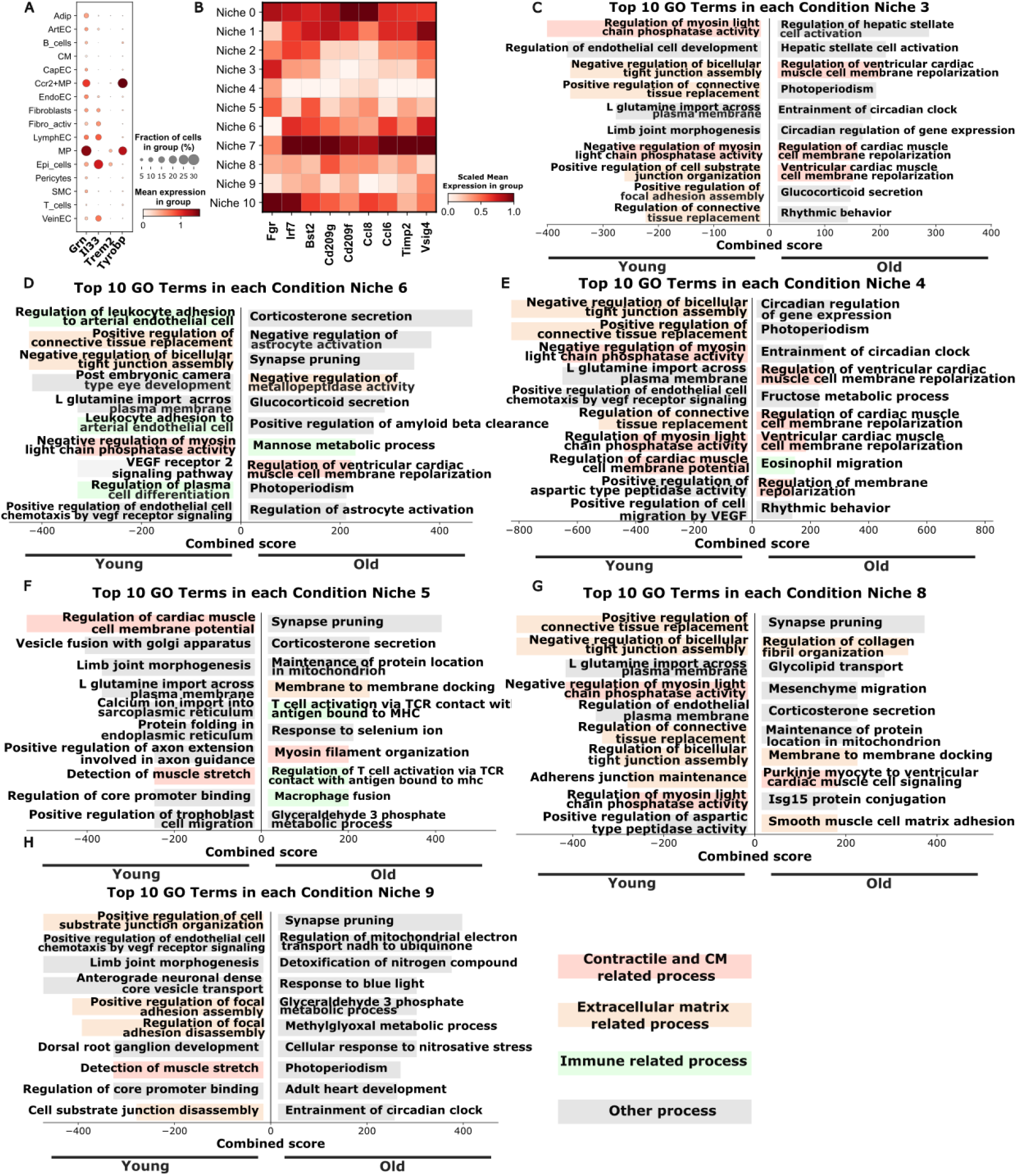
Functional analysis of the cardiac niches. A) Mean expression of genes annotated with the microglia cell activation gene ontology term. B) Mean expression of genes associated with a pro-inflammatory and pro-fibrotic phenotype of macrophages across niches after scaling to [0, 1] across the different niches. C) to H) show the top gene ontology terms regulated in the old (positive axis) and in the young condition (negative axis) for each cardiac niche (3, 6, 4, 5, 8, and 9). Terms were filtered using an adjusted p-value of 0.05 and ranked base on the combined score estimated by enrichR. Terms associated with similar processes were grouped and colored together.

**Extended Figure 6.**
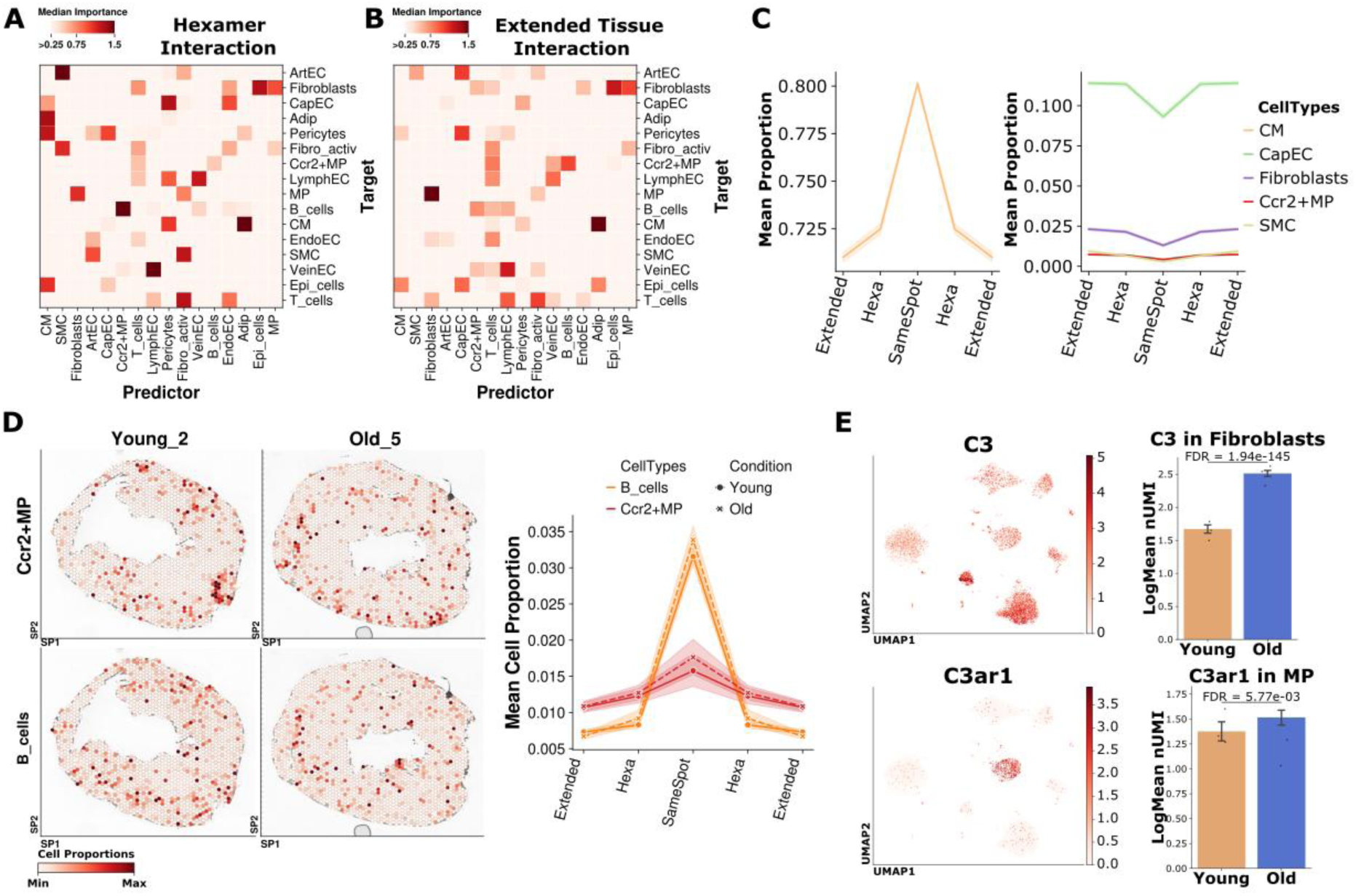
Cell type interactions in the heart. A) and B) Median importance over all samples of one cell type to predict the presence or absence of other cell types in A) the hexamer space (i.e., direct neighboring spots) and B) the extended tissue space (i.e., spots that are not direct neighboring spots). C) Mean proportion of several cell types in the 8 % of spots with the highest abundance of cardiomyocytes (CM). The proportion in the hexamer space (Hexa) and extended tissue (Extended) was mirrored. D) Spatial distribution of the proportion of B cells and Ccr2+MP (right panel) and the mean proportion abundance in the top 8 % of spots with the highest abundance of B cells (left panel). E) UMAP (left panel) and bar plots of the mean expression of *C3* in fibroblasts and *C3ar1* in macrophages (MP) (right value). The adjusted p-value was estimated with Wilcoxon test and Benjamini-Hochberg correction. Only the dataset derived from Vidal et al. was considered. The error bar represents the 95 % confidence interval of the mean.

**Extended Figure 7.**
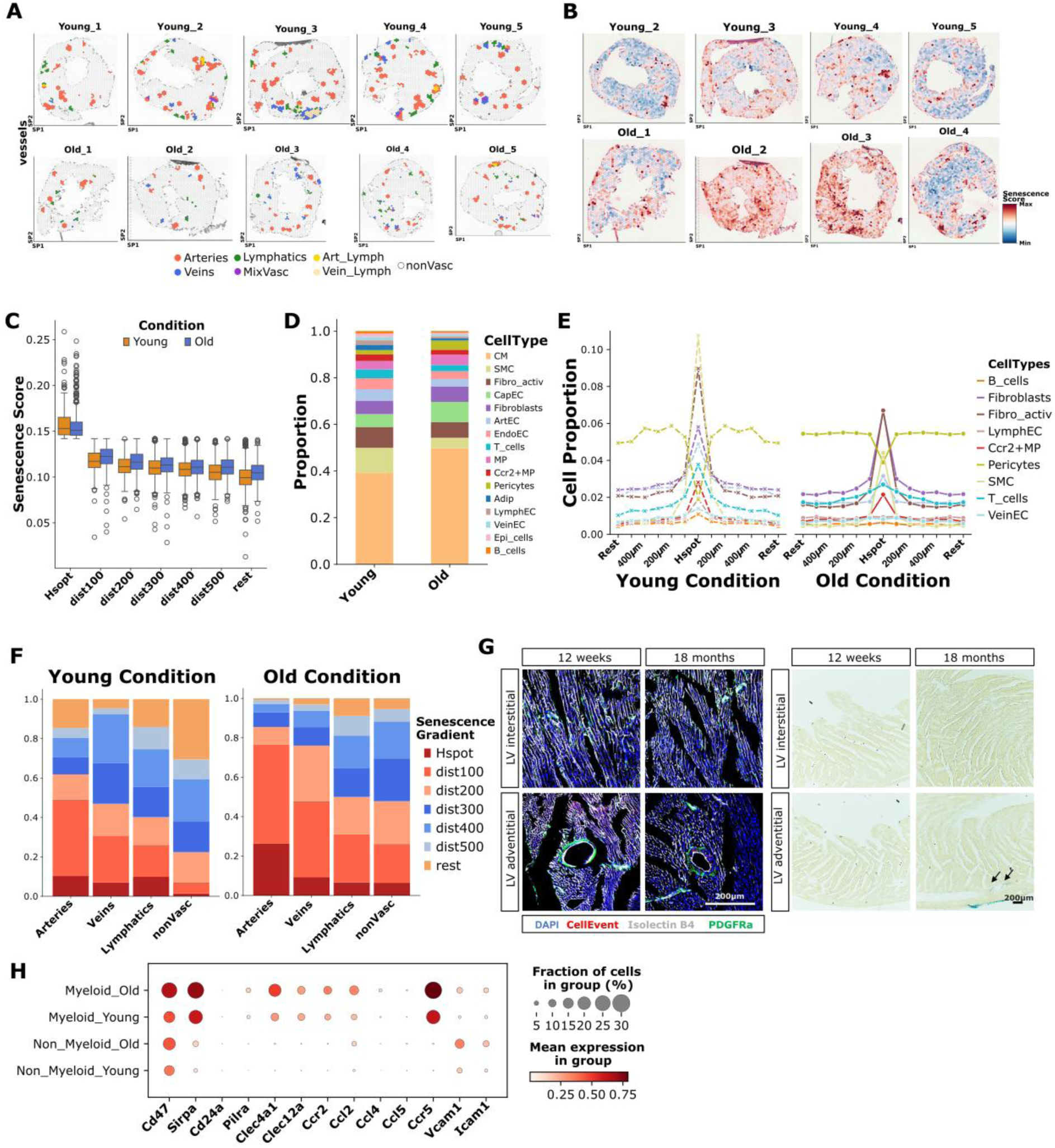
Characterization of cellular senescence during aging in the heart. A) and B) Spatial distribution of A) the vasculature annotation and B) the senescence score. C) Distribution of senescence score in the hotspots of senescence (Hspot) and spots at different distances from the Hspot (in µm) in each condition. D) Proportion of all the cell types across all the Hspot per condition. E) Proportion of different cell types across different locations per condition. The proportion at the different locations was mirrored. F) Proportion of Hspot and spots at different locations (in µm) across the subtype of vessels. G) SA-ß-galactosidase staining and immunofluorescence showing senescence in an interstitial and adventitial region in young and old male mice hearts at low magnification. Isolectin B4 (white, endothelial cells), PDGFRa (green, fibroblasts), CellEvent (red, senescence) and DAPI (nucleus). Scale bars, 200 µm. The arrows in the SA-ß-galactosidase staining indicate regions positive for senescence. H) Mean expression of genes associated to immune inhibitory molecules and immune recruitment in myeloid and non-myeloid cells. Only cells derived from the Vidal et al. dataset were considered.

## References

1. Tsao CW, Aday AW, Almarzooq ZI, Anderson CAM, Arora P, Avery CL, Baker-Smith CM, Beaton AZ, Boehme AK, Buxton AE, et al. Heart Disease and Stroke Statistics-2023 Update: A Report From the American Heart Association. Circulation [Internet]. 2023 [cited 2024 Sep 16];147:E93–E621. Available from: https://pubmed.ncbi.nlm.nih.gov/36695182/

2. Liberale L, Badimon L, Montecucco F, Lüscher TF, Libby P, Camici GG. Inflammation, Aging, and Cardiovascular Disease: JACC Review Topic of the Week. J Am Coll Cardiol [Internet]. 2022 [cited 2025 Feb 24];79:837–847. Available from: https://pubmed.ncbi.nlm.nih.gov/35210039/

3. Litviňuková M, Talavera-López C, Maatz H, Reichart D, Worth CL, Lindberg EL, Kanda M, Polanski K, Heinig M, Lee M, et al. Cells of the adult human heart. Nature 2020 588:7838 [Internet]. 2020 [cited 2025 Feb 8];588:466–472. Available from: https://www.nature.com/articles/s41586-020-2797-4

4. Miranda AMA, Janbandhu V, Maatz H, Kanemaru K, Cranley J, Teichmann SA, Hübner N, Schneider MD, Harvey RP, Noseda M. Single-cell transcriptomics for the assessment of cardiac disease. Nature Reviews Cardiology 2022 20:5 [Internet]. 2022 [cited 2025 Feb 8];20:289–308. Available from: https://www.nature.com/articles/s41569-022-00805-7

5. Cheng M, Jiang Y, Xu J, Mentis AFA, Wang S, Zheng H, Sahu SK, Liu L, Xu X. Spatially resolved transcriptomics: a comprehensive review of their technological advances, applications, and challenges. Journal of Genetics and Genomics. 2023;50:625–640.

6. Farah EN, Hu RK, Kern C, Zhang Q, Lu TY, Ma Q, Tran S, Zhang B, Carlin D, Monell A, et al. Spatially organized cellular communities form the developing human heart. Nature 2024 627:8005 [Internet]. 2024 [cited 2024 Sep 16];627:854–864. Available from: https://www.nature.com/articles/s41586-024-07171-z

7. Kanemaru K, Cranley J, Muraro D, Miranda AMA, Ho SY, Wilbrey-Clark A, Patrick Pett J, Polanski K, Richardson L, Litvinukova M, et al. Spatially resolved multiomics of human cardiac niches. Nature 2023 619:7971 [Internet]. 2023 [cited 2024 Sep 16];619:801–810. Available from: https://www.nature.com/articles/s41586-023-06311-1

8. Wolf FA, Angerer P, Theis FJ. SCANPY: Large-scale single-cell gene expression data analysis. Genome Biol [Internet]. 2018 [cited 2024 Aug 5];19:1–5. Available from: https://genomebiology.biomedcentral.com/articles/10.1186/s13059-017-1382-0

9. Hao Y, Stuart T, Kowalski MH, Choudhary S, Hoffman P, Hartman A, Srivastava A, Molla G, Madad S, Fernandez-Granda C, et al. Dictionary learning for integrative, multimodal and scalable single-cell analysis. Nature Biotechnology 2023 42:2 [Internet]. 2023 [cited 2024 Aug 5];42:293–304. Available from: https://www.nature.com/articles/s41587-023-01767-y

10. Vidal R, Wagner JUG, Braeuning C, Fischer C, Patrick R, Tombor L, Muhly-Reinholz M, John D, Kliem M, Conrad T, et al. Transcriptional heterogeneity of fibroblasts is a hallmark of the aging heart. JCI Insight [Internet]. 2019 [cited 2024 Aug 8];4. Available from: /pmc/articles/PMC6948853/

11. Wagner JUG, Tombor LS, Malacarne PF, Kettenhausen LM, Panthel J, Kujundzic H, Manickam N, Schmitz K, Cipca M, Stilz KA, et al. Aging impairs the neurovascular interface in the heart. Science (1979) [Internet]. 2023 [cited 2024 Aug 8];381:897–906. Available from: https://www.science.org/doi/10.1126/science.ade4961

12. Wolock SL, Lopez R, Klein AM. Scrublet: Computational Identification of Cell Doublets in Single-Cell Transcriptomic Data. Cell Syst. 2019;8:281–291.e9.

13. Lopez R, Regier J, Cole MB, Jordan MI, Yosef N. Deep generative modeling for single-cell transcriptomics. Nature Methods 2018 15:12 [Internet]. 2018 [cited 2024 Aug 5];15:1053–1058. Available from: https://www.nature.com/articles/s41592-018-0229-2

14. Polański K, Young MD, Miao Z, Meyer KB, Teichmann SA, Park JE. BBKNN: fast batch alignment of single cell transcriptomes. Bioinformatics [Internet]. 2020 [cited 2024 Aug 5];36:964–965. Available from: 10.1093/bioinformatics/btz625

15. Xu C, Prete M, Webb S, Jardine L, Stewart BJ, Hoo R, He P, Meyer KB, Teichmann SA. Automatic cell-type harmonization and integration across Human Cell Atlas datasets. Cell [Internet]. 2023 [cited 2024 Aug 5];186:5876-5891.e20. Available from: http://www.cell.com/article/S0092867423013120/fulltext

16. Skelly DA, Squiers GT, McLellan MA, Bolisetty MT, Robson P, Rosenthal NA, Pinto AR. Single-Cell Transcriptional Profiling Reveals Cellular Diversity and Intercommunication in the Mouse Heart. Cell Rep. 2018;22:600–610.

17. Kalucka J, de Rooij LPMH, Goveia J, Rohlenova K, Dumas SJ, Meta E, Conchinha N V., Taverna F, Teuwen LA, Veys K, et al. Single-Cell Transcriptome Atlas of Murine Endothelial Cells. Cell. 2020;180:764–779.e20.

18. Zhou X, Dong K, Zhang S. Integrating spatial transcriptomics data across different conditions, technologies and developmental stages. Nature Computational Science 2023 3:10 [Internet]. 2023 [cited 2024 Aug 6];3:894–906. Available from: https://www.nature.com/articles/s43588-023-00528-w

19. Badia-I-Mompel P, Vélez Santiago J, Braunger J, Geiss C, Dimitrov D, Müller-Dott S, Taus P, Dugourd A, Holland CH, Ramirez Flores RO, et al. decoupleR: ensemble of computational methods to infer biological activities from omics data. Bioinformatics Advances [Internet]. 2022 [cited 2024 Sep 19];2. Available from: 10.1093/bioadv/vbac016

20. Kleshchevnikov V, Shmatko A, Dann E, Aivazidis A, King HW, Li T, Elmentaite R, Lomakin A, Kedlian V, Gayoso A, et al. Cell2location maps fine-grained cell types in spatial transcriptomics. Nature Biotechnology 2022 40:5 [Internet]. 2022 [cited 2024 Aug 6];40:661–671. Available from: https://www.nature.com/articles/s41587-021-01139-4

21. Schubert M, Klinger B, Klünemann M, Sieber A, Uhlitz F, Sauer S, Garnett MJ, Blüthgen N, Saez-Rodriguez J. Perturbation-response genes reveal signaling footprints in cancer gene expression. Nat Commun [Internet]. 2018 [cited 2024 Sep 4];9:20–20. Available from: https://europepmc.org/articles/PMC5750219

22. Liberzon A, Subramanian A, Pinchback R, Thorvaldsdóttir H, Tamayo P, Mesirov JP. Molecular signatures database (MSigDB) 3.0. Bioinformatics [Internet]. 2011 [cited 2024 Nov 27];27:1739–1740. Available from: 10.1093/bioinformatics/btr260

23. Castanza AS, Recla JM, Eby D, Thorvaldsdóttir H, Bult CJ, Mesirov JP. Extending support for mouse data in the Molecular Signatures Database (MSigDB). Nature Methods 2023 20:11 [Internet]. 2023 [cited 2024 Nov 27];20:1619–1620. Available from: https://www.nature.com/articles/s41592-023-02014-7

24. Love MI, Huber W, Anders S. Moderated estimation of fold change and dispersion for RNA-seq data with DESeq2. Genome Biol [Internet]. 2014 [cited 2024 Nov 28];15:550. Available from: https://pmc.ncbi.nlm.nih.gov/articles/PMC4302049/

25. Tanevski J, Flores ROR, Gabor A, Schapiro D, Saez-Rodriguez J. Explainable multiview framework for dissecting spatial relationships from highly multiplexed data. Genome Biol [Internet]. 2022 [cited 2024 Sep 18];23:1–31. Available from: https://genomebiology.biomedcentral.com/articles/10.1186/s13059-022-02663-5

26. Li H, Ma T, Hao M, Guo W, Gu J, Zhang X, Wei L. Decoding functional cell-cell communication events by multi-view graph learning on spatial transcriptomics. Brief Bioinform [Internet]. 2023 [cited 2024 Nov 21];24. Available from: https://pubmed.ncbi.nlm.nih.gov/37824741/

27. Tirosh I, Izar B, Prakadan SM, Wadsworth MH, Treacy D, Trombetta JJ, Rotem A, Rodman C, Lian C, Murphy G, et al. Dissecting the multicellular ecosystem of metastatic melanoma by single-cell RNA-seq. Science [Internet]. 2016 [cited 2024 Nov 28];352:189. Available from: https://pmc.ncbi.nlm.nih.gov/articles/PMC4944528/

28. Satija R, Farrell JA, Gennert D, Schier AF, Regev A. Spatial reconstruction of single-cell gene expression data. Nature Biotechnology 2015 33:5 [Internet]. 2015 [cited 2024 Nov 28];33:495–502. Available from: https://www.nature.com/articles/nbt.3192

29. Avelar RA, Ortega JG, Tacutu R, Tyler EJ, Bennett D, Binetti P, Budovsky A, Chatsirisupachai K, Johnson E, Murray A, et al. A multidimensional systems biology analysis of cellular senescence in aging and disease. Genome Biol [Internet]. 2020 [cited 2024 Nov 27];21. Available from: https://pubmed.ncbi.nlm.nih.gov/32264951/

30. Liu GH, Bao Y, Qu J, Zhang W, Zhang T, Kang W, Yang F, Ji Q, Jiang X, Ma Y, et al. Aging Atlas: a multi-omics database for aging biology. Nucleic Acids Res [Internet]. 2021 [cited 2024 Nov 27];49:D825–D830. Available from: https://pubmed.ncbi.nlm.nih.gov/33119753/

31. Saul D, Kosinsky RL, Atkinson EJ, Doolittle ML, Zhang X, LeBrasseur NK, Pignolo RJ, Robbins PD, Niedernhofer LJ, Ikeno Y, et al. A new gene set identifies senescent cells and predicts senescence-associated pathways across tissues. Nature Communications 2022 13:1 [Internet]. 2022 [cited 2024 Sep 17];13:1–15. Available from: https://www.nature.com/articles/s41467-022-32552-1

32. Fridman AL, Tainsky MA. Critical pathways in cellular senescence and immortalization revealed by gene expression profiling. Oncogene 2008 27:46 [Internet]. 2008 [cited 2024 Nov 27];27:5975–5987. Available from: https://www.nature.com/articles/onc2008213

33. Domínguez Conde C, Xu C, Jarvis LB, Rainbow DB, Wells SB, Gomes T, Howlett SK, Suchanek O, Polanski K, King HW, et al. Cross-tissue immune cell analysis reveals tissue-specific features in humans. Science (1979) [Internet]. 2022 [cited 2024 Aug 29];376. Available from: https://www.science.org/doi/10.1126/science.abl5197

34. Hocker JD, Poirion OB, Zhu F, Buchanan J, Zhang K, Chiou J, Wang TM, Zhang Q, Hou X, Li YE, et al. Cardiac cell type-specific gene regulatory programs and disease risk association. Sci Adv [Internet]. 2021 [cited 2024 Sep 16];7. Available from: https://www.science.org/doi/10.1126/sciadv.abf1444

35. Koenig AL, Shchukina I, Amrute J, Andhey PS, Zaitsev K, Lai L, Bajpai G, Bredemeyer A, Smith G, Jones C, et al. Single-cell transcriptomics reveals cell-type-specific diversification in human heart failure. Nature Cardiovascular Research 2022 1:3 [Internet]. 2022 [cited 2024 Sep 16];1:263–280. Available from: https://www.nature.com/articles/s44161-022-00028-6

36. Litviňuková M, Talavera-López C, Maatz H, Reichart D, Worth CL, Lindberg EL, Kanda M, Polanski K, Heinig M, Lee M, et al. Cells of the adult human heart. Nature 2020 588:7838 [Internet]. 2020 [cited 2024 Sep 16];588:466–472. Available from: https://www.nature.com/articles/s41586-020-2797-4

37. Lavine K. Identification of inflammatory lipid-associated macrophages in human carotid atherosclerosis. Nature Cardiovascular Research 2023 2:7 [Internet]. 2023 [cited 2024 Oct 15];2:604–605. Available from: https://www.nature.com/articles/s44161-023-00299-7

38. Su H, Cantrell AC, Zeng H, Zhu SH, Chen JX. Emerging Role of Pericytes and Their Secretome in the Heart. Cells [Internet]. 2021 [cited 2024 Sep 4];10:1–22. Available from: /pmc/articles/PMC8001346/

39. Bernal-Ramirez J, Diaz-Vesga MC, Talamilla M, Mendez A, Quiroga C, Garza-Cervantes JA, Lazaro-Alfaro A, Jerjes-Sanchez C, Henriquez M, Garcia-Rivas G, et al. Exploring Functional Differences between the Right and Left Ventricles to Better Understand Right Ventricular Dysfunction. Oxid Med Cell Longev [Internet]. 2021 [cited 2024 Oct 15];2021. Available from: /pmc/articles/PMC8421158/

40. North BJ, Sinclair DA. The intersection between aging and cardiovascular disease. Circ Res [Internet]. 2012 [cited 2024 Oct 22];110:1097–1108. Available from: https://www.ahajournals.org/doi/10.1161/CIRCRESAHA.111.246876

41. Wagner JUG, Tombor LS, Malacarne PF, Kettenhausen LM, Panthel J, Kujundzic H, Manickam N, Schmitz K, Cipca M, Stilz KA, et al. Aging impairs the neurovascular interface in the heart. Science [Internet]. 2023 [cited 2025 Jan 6];381:897–906. Available from: https://pubmed.ncbi.nlm.nih.gov/37616346/

42. Acín-Pérez R, Iborra S, Martí-Mateos Y, Cook ECL, Conde-Garrosa R, Petcherski A, Muñoz M ^a^M, Martínez de Mena R, Krishnan KC, Jiménez C, et al. Fgr kinase is required for proinflammatory macrophage activation during diet-induced obesity. Nat Metab [Internet]. 2020 [cited 2025 Feb 4];2:974–988. Available from: https://pubmed.ncbi.nlm.nih.gov/32943786/

43. Piedra-Quintero ZL, Serrano C, Villegas-Sepúlveda N, Maravillas-Montero JL, Romero-Ramírez S, Shibayama M, Medina-Contreras O, Nava P, Santos-Argumedo L. Myosin 1F Regulates M1-Polarization by Stimulating Intercellular Adhesion in Macrophages. Front Immunol [Internet]. 2019 [cited 2025 Feb 4];9. Available from: https://pubmed.ncbi.nlm.nih.gov/30687322/

44. Zhang YZ, Wu Y, Li MJ, Mijiti A, Cheng LF. Identification of macrophage driver genes in fibrosis caused by different heart diseases based on omics integration. J Transl Med [Internet]. 2024 [cited 2025 Jan 15];22:839. Available from: https://pmc.ncbi.nlm.nih.gov/articles/PMC11391649/

45. Nagai T, Honda S, Sugano Y, Matsuyama T aki, Ohta-Ogo K, Asaumi Y, Ikeda Y, Kusano K, Ishihara M, Yasuda S, et al. Decreased myocardial dendritic cells is associated with impaired reparative fibrosis and development of cardiac rupture after myocardial infarction in humans. J Am Heart Assoc [Internet]. 2014 [cited 2025 Jan 15];3. Available from: https://www.ahajournals.org/doi/10.1161/JAHA.114.000839

46. Mak TW, Hauck L, Grothe D, Billia F. p53 regulates the cardiac transcriptome. Proc Natl Acad Sci U S A [Internet]. 2017 [cited 2024 Oct 28];114:2331–2336. Available from: https://www.pnas.org/doi/abs/10.1073/pnas.1621436114

47. Thüroff JW, Hort W, Lichti H. Diameter of coronary arteries in 36 species of mammalian from mouse to giraffe. Basic Res Cardiol [Internet]. 1984 [cited 2024 Aug 8];79:199–206. Available from: https://link.springer.com/article/10.1007/BF01908306

48. Müller B, Lang S, Dominietto M, Rudin M, Schulz G, Deyhle H, Germann M, Pfeiffer F, David C, Weitkamp T. High-resolution tomographic imaging of microvessels. 10.1117/12.794157 [Internet]. 2008 [cited 2024 Aug 8];7078:89–98. Available from: https://www.spiedigitallibrary.org/conference-proceedings-of-spie/7078/70780B/High-resolution-tomographic-imaging-of-microvessels/10.1117/12.794157.full

49. Adamo L, Rocha-Resende C, Lin CY, Evans S, Williams J, Dun H, Li W, Mpoy C, Andhey PS, Rogers BE, et al. Myocardial B cells are a subset of circulating lymphocytes with delayed transit through the heart. JCI Insight [Internet]. 2020 [cited 2024 Nov 1];5. Available from: https://pubmed.ncbi.nlm.nih.gov/31945014/

50. Subramanian A, Tamayo P, Mootha VK, Mukherjee S, Ebert BL, Gillette MA, Paulovich A, Pomeroy SL, Golub TR, Lander ES, et al. Gene set enrichment analysis: A knowledge-based approach for interpreting genome-wide expression profiles. Proc Natl Acad Sci U S A [Internet]. 2005 [cited 2024 Nov 27];102:15545–15550. Available from: https://www.pnas.org/doi/abs/10.1073/pnas.0506580102

51. Ji Z, Zhang B, Zhang W, Gu Y, Liu G. Spatial transcriptomic landscape unveils immunoglobin-associated senescence as a hallmark of aging. Cell. 2024;1–20.

52. Radu-Ioniţa F, Bontaş E, Goleanu V, Cîrciumaru B, Bartoş D, Parepa I, Ţintoiu IC, Popa A. Heart Embryology: Overview. Right Heart Pathology: From Mechanism to Management [Internet]. 2018 [cited 2025 Feb 13];3–24. Available from: https://link.springer.com/chapter/10.1007/978-3-319-73764-5_1

53. Cadosch N, Gil-Cruz C, Perez-Shibayama C, Ludewig B. Cardiac Fibroblastic Niches in Homeostasis and Inflammation. Circ Res [Internet]. 2024 [cited 2025 Feb 13];134:1703–1717. Available from: https://pubmed.ncbi.nlm.nih.gov/38843287/

54. Torp MK, Ranheim T, Schjalm C, Hjorth M, Heiestad CM, Dalen KT, Nilsson PH, Mollnes TE, Pischke SE, Lien E, et al. Intracellular Complement Component 3 Attenuated Ischemia-Reperfusion Injury in the Isolated Buffer-Perfused Mouse Heart and Is Associated With Improved Metabolic Homeostasis. Front Immunol [Internet]. 2022 [cited 2025 Feb 17];13. Available from: https://pubmed.ncbi.nlm.nih.gov/35432387/

55. Ito S, Hashimoto H, Yamakawa H, Kusumoto D, Akiba Y, Nakamura T, Momoi M, Komuro J, Katsuki T, Kimura M, et al. The complement C3-complement factor D-C3a receptor signalling axis regulates cardiac remodelling in right ventricular failure. Nat Commun [Internet]. 2022 [cited 2025 Feb 17];13. Available from: https://pubmed.ncbi.nlm.nih.gov/36109509/

56. Wei LL, Ma N, Wu KY, Wang JX, Diao TY, Zhao SJ, Bai L, Liu E, Li ZF, Zhou W, et al. Protective Role of C3aR (C3a Anaphylatoxin Receptor) Against Atherosclerosis in Atherosclerosis-Prone Mice. Arterioscler Thromb Vasc Biol [Internet]. 2020 [cited 2025 Feb 17];40:2070–2083. Available from: https://pubmed.ncbi.nlm.nih.gov/32762445/

57. Fu L, Yin C, Zhao Q, Guo S, Shao W, Xia T, Sun Q, Chen L, Wang M, Xia H. CD81+ fibroblasts, a unique subpopulation with accelerated cellular senescence, exaggerate inflammation and activate neutrophils via C3/C3aR1 axis in periodontitis. Elife [Internet]. 2024 [cited 2025 Feb 17];13. Available from: https://elifesciences.org/reviewed-preprints/96908

58. Ruiz-Orera J, Miller DC, Greiner J, Genehr C, Grammatikaki A, Blachut S, Mbebi J, Patone G, Myronova A, Adami E, et al. Evolution of translational control and the emergence of genes and open reading frames in human and non-human primate hearts. Nature Cardiovascular Research 2024 3:10 [Internet]. 2024 [cited 2025 Feb 17];3:1217–1235. Available from: https://www.nature.com/articles/s44161-024-00544-7

59. Sabbatinelli J, Prattichizzo F, Olivieri F, Procopio AD, Rippo MR, Giuliani A. Where Metabolism Meets Senescence: Focus on Endothelial Cells. Front Physiol [Internet]. 2019 [cited 2024 Oct 22];10:1523. Available from: https://pmc.ncbi.nlm.nih.gov/articles/PMC6930181/

60. Wiley CD, Campisi J. The metabolic roots of senescence: mechanisms and opportunities for intervention. Nat Metab [Internet]. 2021 [cited 2024 Oct 22];3:1290. Available from: https://pmc.ncbi.nlm.nih.gov/articles/PMC8889622/

61. Elder SS, Emmerson E. Senescent cells and macrophages: key players for regeneration? Open Biol [Internet]. 2020 [cited 2025 Feb 13];10:200309. Available from: https://pmc.ncbi.nlm.nih.gov/articles/PMC7776574/

62. Deng H, Wang G, Zhao S, Tao Y, Zhang Z, Yang J, Lei Y. New hope for tumor immunotherapy: the macrophage-related “do not eat me” signaling pathway. Front Pharmacol [Internet]. 2023 [cited 2025 Feb 13];14:1228962. Available from: https://pubmed.ncbi.nlm.nih.gov/37484024/

63. Uto T, Fukaya T, Takagi H, Arimura K, Nakamura T, Kojima N, Malissen B, Sato K. Clec4A4 is a regulatory receptor for dendritic cells that impairs inflammation and T-cell immunity. Nature Communications 2016 7:1 [Internet]. 2016 [cited 2025 Feb 14];7:1–15. Available from: https://www.nature.com/articles/ncomms11273

64. Li Q, Yang Z, He X, Yang X. Comprehensive analysis of PILRΑ’s association with the prognosis, tumor immune infiltration, and immunotherapy in pan-cancer. Scientific Reports 2023 13:1 [Internet]. 2023 [cited 2025 Feb 14];13:1–18. Available from: https://www.nature.com/articles/s41598-023-41649-6

65. Gracia-Hernandez M, Suresh M, Villagra A. The advances in targeting CD47/SIRPα “do not eat me” axis and their ongoing challenges as an anticancer therapy. Oncotarget [Internet]. 2024 [cited 2025 Feb 14];15:462–465. Available from: https://pubmed.ncbi.nlm.nih.gov/38985136/

66. Lecacheur M, Ammerlaan DJM, Dierickx P. Circadian rhythms in cardiovascular (dys)function: approaches for future therapeutics. npj Cardiovascular Health 2024 1:1 [Internet]. 2024 [cited 2025 Jan 15];1:1–13. Available from: https://www.nature.com/articles/s44325-024-00024-8

67. Hayter EA, Wehrens SMT, Van Dongen HPA, Stangherlin A, Gaddameedhi S, Crooks E, Barron NJ, Venetucci LA, Neill JSO, Brown TM, et al. Distinct circadian mechanisms govern cardiac rhythms and susceptibility to arrhythmia. Nature Communications 2021 12:1 [Internet]. 2021 [cited 2025 Jan 15];12:1–13. Available from: https://www.nature.com/articles/s41467-021-22788-8

68. Levine B, Kalman J, Mayer L, Fillit HM, Packer M. Elevated Circulating Levels of Tumor Necrosis Factor in Severe Chronic Heart Failure. New England Journal of Medicine [Internet]. 1990 [cited 2025 Feb 14];323:236–241. Available from: https://www.nejm.org/doi/full/10.1056/NEJM199007263230405

69. Baumgarten G, Knuefermann P, Mann DL. Cytokines as emerging targets in the treatment of heart failure. Trends Cardiovasc Med [Internet]. 2000 [cited 2025 Feb 14];10:216–223. Available from: https://pubmed.ncbi.nlm.nih.gov/11282298/

70. Horitani K, Shiojima I. Wnt signaling in cardiac development and heart diseases. In Vitro Cell Dev Biol Anim [Internet]. 2024 [cited 2025 Feb 14];60:482. Available from: https://pmc.ncbi.nlm.nih.gov/articles/PMC11126472/

71. Mak TW, Hauck L, Grothe D, Billia F. p53 regulates the cardiac transcriptome. Proc Natl Acad Sci U S A [Internet]. 2017 [cited 2024 Dec 12];114:2331–2336. Available from: https://pmc.ncbi.nlm.nih.gov/articles/PMC5338492/

72. Rastogi C, Rube HT, Kribelbauer JF, Crocker J, Loker RE, Martini GD, Laptenko O, Freed-Pastor WA, Prives C, Stern DL, et al. Counteracting age-related VEGF signaling insufficiency promotes healthy aging and extends life span. Science (1979) [Internet]. 2021 [cited 2025 Feb 17];373:E3692–E3701. Available from: https://www.science.org/doi/10.1126/science.abc8479

